# Quinone extraction drives atmospheric carbon monoxide oxidation in bacteria

**DOI:** 10.1101/2024.01.09.574806

**Authors:** Ashleigh Kropp, David L. Gillett, Hari Venugopal, Miguel A. Gonzálvez, James P. Lingford, Christopher K. Barlow, Jie Zhang, Chris Greening, Rhys Grinter

## Abstract

Diverse bacteria and archaea use atmospheric carbon monoxide (CO) as an energy source during long-term survival. This process enhances the biodiversity of soil and marine ecosystems globally and removes 250 million tonnes of a toxic, climate-relevant pollutant from the atmosphere each year. Bacteria use [MoCu]-carbon monoxide dehydrogenases (Mo-CODH) to convert CO to carbon dioxide, then transfer the liberated high-energy electrons to the aerobic respiratory chain. However, given no high-affinity Mo-CODH has been purified, it is unknown how these enzymes oxidise CO at low concentrations and interact with the respiratory chain. Here we resolve these knowledge gaps by analysing Mo-CODH (CoxSML) and its hypothetical partner CoxG from *Mycobacterium smegmatis*. Kinetic and electrochemical analyses show purified Mo-CODH is a highly active high-affinity enzyme (*K*_m_ = 139 nM, *k*_cat_ = 54.2 s^-1^). Based on its 1.85 Å resolution cryoEM structure, Mo-CODH forms a CoxSML homodimer similar to characterised low-affinity homologs, but has distinct active site coordination and narrower gas channels that may modulate affinity. We provide structural, biochemical, and genetic evidence that Mo-CODH transfers CO-derived electrons to the aerobic respiratory chain *via* the membrane-bound menaquinone-binding protein CoxG. Consistently, CoxG is required for CO-driven respiration, extracts menaquinone from mycobacterial membranes, and binds quinones in a hydrophobic pocket. Finally, we show that Mo-CODH and CoxG genetically and structurally associate in diverse bacteria and archaea. These findings reveal the basis of a biogeochemically and ecologically important process, while demonstrating that the newly discovered process of long-range quinone transport is a general mechanism of energy conservation, which convergently evolved on multiple occasions.

## Introduction

Carbon monoxide (CO) is both a potent poison of multicellular life and a high-energy fuel and carbon source for microorganisms ^1–5^. CO is released into the atmosphere in vast quantities, with natural and anthropogenic sources contributing an estimated 2,600 million tonnes of CO emissions annually ^6,7^. Despite this, average CO concentrations in the atmosphere remain extremely low, at around 100 parts per billion (ppbv), due to consumption by abiotic processes and microbial oxidation ^6,8^. Microbial consumption accounts for an estimated 10-15% of CO removed from the atmosphere (approximately 250 million tonnes annually) ^7^. This biogeochemically important process is mediated by microbes that encode high-affinity variants of the form I molybdenum-copper dependent carbon monoxide dehydrogenases (Mo-CODH). In microbial cells, these enzymes oxidise CO at atmospheric concentrations, and the high-energy electrons derived from this process are transferred to the aerobic respiratory chain ^4,6,9–11^. Due to its low concentration, the energy provided by oxidising atmospheric CO is generally insufficient to support growth and replication ^6,12^. However, CO provides a supplemental energy source for vast numbers of dormant bacteria present in soils, oceans, and other nutrient-deprived environments, allowing them to persist during starvation ^4,6,13–15^. The ability to oxidise atmospheric CO is widespread, with bacteria and archaea from 17 phyla encoding a Mo-CODH predicted to be capable of mediating this process, and isolates from four phyla have been shown experimentally to oxidise atmospheric CO ^6^.

Despite the biogeochemical and ecological importance of microbial atmospheric CO oxidation, no Mo-CODH has been isolated that has been shown to be capable of mediating this process. Mo-CODHs have been isolated and structurally and biochemically characterised from aerobic carboxydotrophic bacteria, which use high concentrations of CO as both a carbon and energy source for autotrophic growth ^16–21^. However, these enzymes have a lower affinity for CO (K_m_ > 400 nM) which is incompatible with oxidation of the gas at atmospheric concentrations ^4,22–25^. Through studies on whole cells, we recently demonstrated that a Mo-CODH encoded by *Mycobacterium smegmatis* (Mo-CODH_Ms_) enables the bacterium to oxidise atmospheric CO, which enhances its survival during starvation ^4^. We also showed that electrons derived from Mo-CODH_Ms_ are transferred to terminal oxidases and thus support aerobic respiration ^4,5^. However, the structural and biochemical basis for the high-affinity kinetics of Mo-CODH_Ms_ remains unknown.

Also unresolved are the physical associations and electron pathways made by Mo-CODH. Considering Mo-CODH is a respiratory enzyme, it must either directly or indirectly reduce membrane-bound quinone ^4^. Despite a lack of integral or peripheral membrane-associating regions, Mo-CODH from *Afipia carboxidovorans* (Mo-CODH_Ac_) is predominantly membrane-associated ^26^. In contrast, more limited membrane association is observed with Mo-CODH_Ms_, which may reflect the differing roles of Mo-CODH in these organisms, in growth and persistence respectively ^4,18,26,27^. Previous reports indicate that Mo-CODH_Ac_ reduces respiratory quinone analogues, suggesting that quinones directly accept electrons from this enzyme ^28^. However, it remains unclear how hydrophobic quinones reach the electron acceptor site of these soluble enzymes. It has been suggested that the Mo-CODH-associated protein CoxG plays a role in this process ^29^. In *A. carboxidovorans,* CoxG is not required for Mo-CODH_Ac_ assembly or maturation, but it is essential for mediating membrane association ^18,26^. The loss of CoxG does not abolish CO oxidation by Mo-CODH_Ac_, but greatly increases generation time (∼7-fold) when CO is the sole substrate for growth ^29^. These observations led to the hypothesis that CoxG recruits soluble CO dehydrogenase to the cytoplasmic membrane, thus enabling electron transfer to quinones in the membrane ^29^.

In this study, we aimed to resolve the structural and biochemical basis for the high-affinity of Mo-CODH_Ms_, and its interactions with the electron transport chain. To do so, we isolated both Mo-CODH_Ms_ and CoxG, and integrated cryoEM, X-ray crystallography, kinetic, electrochemical, and genetic analyses to show the structural basis of atmospheric CO oxidation and demonstrate how electrons are transferred to the respiratory chain.

## Main

### Mo-CODH from *M. smegmatis* oxidises CO at atmospheric concentrations

We isolated Mo-CODH_Ms_ using a chromosomally encoded Strep-tag II on its CoxM subunit. Due to the complex maturation of this enzyme family, which requires at least four genetically associated assembly factors ^29,30^, Mo-CODH_Ms_ was expressed chromosomally at native levels, with cells harvested at late stationary phase when the enzyme is most highly expressed ^4^. Though yield was low (∼2-5 µg of enzyme per litre of *M. smegmatis* culture), this strategy yielded high-purity protein, with SDS-PAGE analysis indicating that only the expected CoxS (17 kDa), CoxM (35 kDa) and CoxL (86 kDa) subunits were present in the complex (Figure S1a,b). Mo-CODH is active and can reduce menadione, a soluble analogue of the membrane-bound electron carrier menaquinone, as an electron acceptor (Figure 1a).

**Figure 1.**
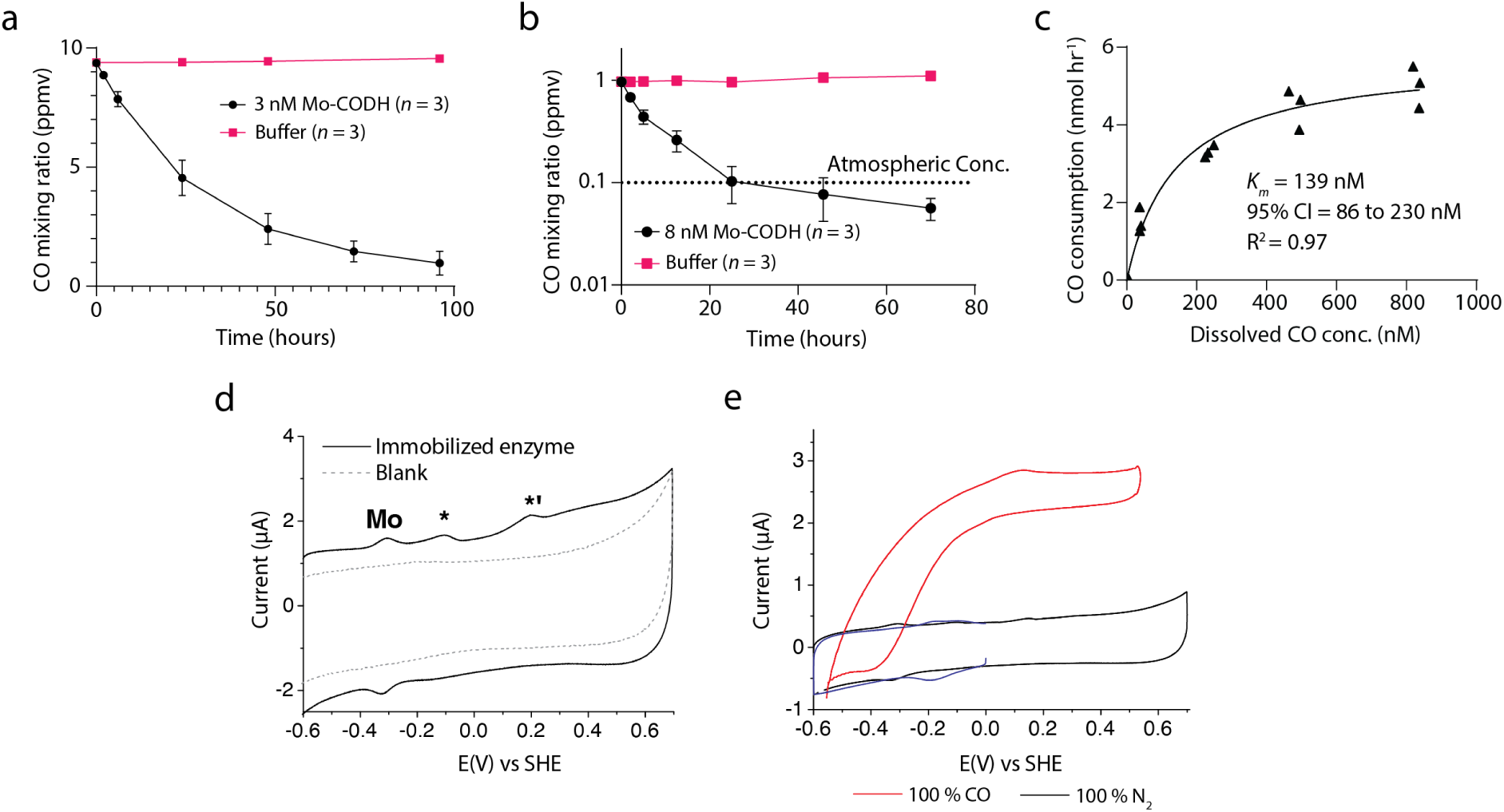
Mo-CODH_Ms_ oxidises CO with high catalytic efficiency to below atmospheric concentrations. **(a)** Gas chromatography analysis of the CO concentration of the headspace of sealed vials containing Mo-CODH_Ms_ or no enzyme, with 200 μM **menadione** as the electron acceptor. **(b)** Gas chromatography analysis of the CO concentration of the headspace of sealed vials containing Mo-CODH_Ms_ or no enzyme, with 50 μM **methylene blue** as the electron acceptor. Mo-CODH_Ms_ can oxidise CO below atmospheric concentrations (black dotted line). In **a** and **b**, data are mean ± s.d. **(c)** Steady-state Michaelis–Menten kinetics of Mo-CODH_Ms_ consumption CO in the headspace of sealed vials with 50 μM methylene blue as the electron acceptor. **(d)** DC voltammogram of surface-confined Mo-CODH_Ms_ on a PIGE electrode obtained at pH 8.0, under N_2_ (scan rate = 500 mV s^−1^, *T* = 21 °C). **(e)** Comparison of the DC voltammograms obtained at pH 8.0, under N_2_-(black line), CO-(red), and CO_2_-saturated (blue) conditions (scan rate = 50 mV s^−1^, *T* = 21 °C).

Although Mo-CODH_Ms_ can oxidise atmospheric CO in whole cells and provide the resulting electrons to the respiratory electron transport chain, other Mo-CODHs that have been isolated are low-affinity enzymes likely incapable of oxidising CO at this concentration ^23–25^. As such, it was unclear whether the *M. smegmatis* enzyme has an inherently high affinity for CO or if this property results from coupling to the respiratory electron transport chain. To resolve this question, we tested the ability of purified Mo-CODH_Ms_ to oxidise 1 part per million (ppm) CO in an ambient air headspace using gas chromatography. Mo-CODH_Ms_ consumed CO below atmospheric levels (to 0.065 ppm in headspace, corresponding to 51 pM in solution) using the artificial electron acceptor methylene blue (Figure 1b). To determine the affinity of Mo-CODH_Ms_ for CO oxidation, we performed kinetic analysis by measuring headspace CO consumption at different concentrations. These data were fitted with Michaelis-Menten kinetics, allowing us to calculate an apparent *K_m_* of 139 nM (95% CI 86 to 230 nM) for CO oxidation by Mo-CODH_Ms_ (Figure 1c), which is significantly lower than that previously reported for a Mo-CODH and consistent with measurements of CO affinity of mycobacterial whole cells ^4,23–25^.

To investigate the electrochemical properties of Mo-CODH_Ms_, we performed protein film electrochemistry (PFE) on the purified enzyme. Cyclic voltammograms on a film of Mo-CODH_Ms_ in 100% N_2_ at pH 8.0 revealed three oxidation peaks that likely correspond to redox transitions of the enzymes Mo (−0.305 V vs SHE, pH 8.0), FeS (−0.120 V) and FAD (+0.185 V) cofactors (Figure 1d; supplemental note 1). A positive sigmoidal shaped steady-state current was observed in voltammograms in the presence of CO, corresponding to oxidation of the gas, with an onset potential close to the equilibrium potential of the CO_2_/CO reaction (−0.58 V vs SHE at pH 8.0^31^), indicating that Mo-CODH_Ms_ operates without a significant overpotential (Figure 1e). By calculating Mo-CODH_Ms_ loading, we were able to derive a turnover frequency (*k_cat_*) of 54.2 s^-1^ for the enzyme, which is comparable to 93.3 s^-1^ determined previously for Mo-CODH_Ac_ (Figure S2; supplemental note 1) ^32^. Based on *k*_cat_/*K*_m_, Mo-CODH_Ms_ is an extremely efficient enzyme (3.9 × 10^8^ M^-1^ s^-1^) approaching theoretical diffusion limits. Based on this analysis, Mo-CODH_Ms_ exhibits similar redox properties compared to Mo-CODH from other organisms, and may derive its high affinity from structural modifications.

### Mo-CODH_Ms_ exhibits novel active site cofactor coordination and narrow gas channels

To determine the structure of Mo-CODH_Ms_ we performed cryo-electron microscopy (cryoEM) imaging and single-particle reconstruction of the purified 272 kDa complex. CryoEM micrographs and resulting class averages revealed a *C_2_* symmetrical molecule, which is consistent with the previously determined structures of low-affinity Mo-CODHs (Figure S3a, b) ^16,22,33^. Despite indications that Mo-CODH utilises membrane-localised quinones as an electron acceptor (Figure 1a) ^28^, Mo-CODH_Ms_ did not co-purify with membranes nor were any additional electron transferring subunits identified in class averages. Data processing and 3D reconstruction yielded maps with a nominal resolution of 1.85 Å, which allowed us to build and refine the structure of Mo-CODH_Ms_ with a high degree of precision (Figure 2a; Figure S3c-g; Table S1; Movie S1; Movie S2). Like other enzymes of this family, Mo-CODH_Ms_ is a heterohexamer composed of two CoxSML subunits that interact via their CoxL subunits (Figure 2a). The CoxL subunit encloses a CuMo-molybdopterin cytosine dinucleotide (CuMCD) containing active site that oxidises CO and transfers the resulting electrons to a flavin adenosine dinucleotide (FAD) cofactor at the electron acceptor site in CoxM, via two [2Fe-2S] clusters in the CoxS subunit (Figure 2b). Despite forming a dimer, the electron transfer pathways of the two CoxSML subunits are not within electron transfer distance (Figure S4a). Mo-CODH mediates catalysis by coordinating CO between the Cu and Mo ions of the active site ^34–36^. The bond lengths in this region of the cofactor indicate that Mo-CODH_Ms_ is in a reduced state, with the Mo ion coordinated by oxo and hydroxyl ligand, in addition to the two thiol groups of the pterin, and the Cu bridging sulphur (Figure 2c) ^16,23,37^. This reduced state may result from the oxidation of atmospheric CO, reducing the cofactor in the absence of an exogenous electron acceptor.

**Figure 2.**
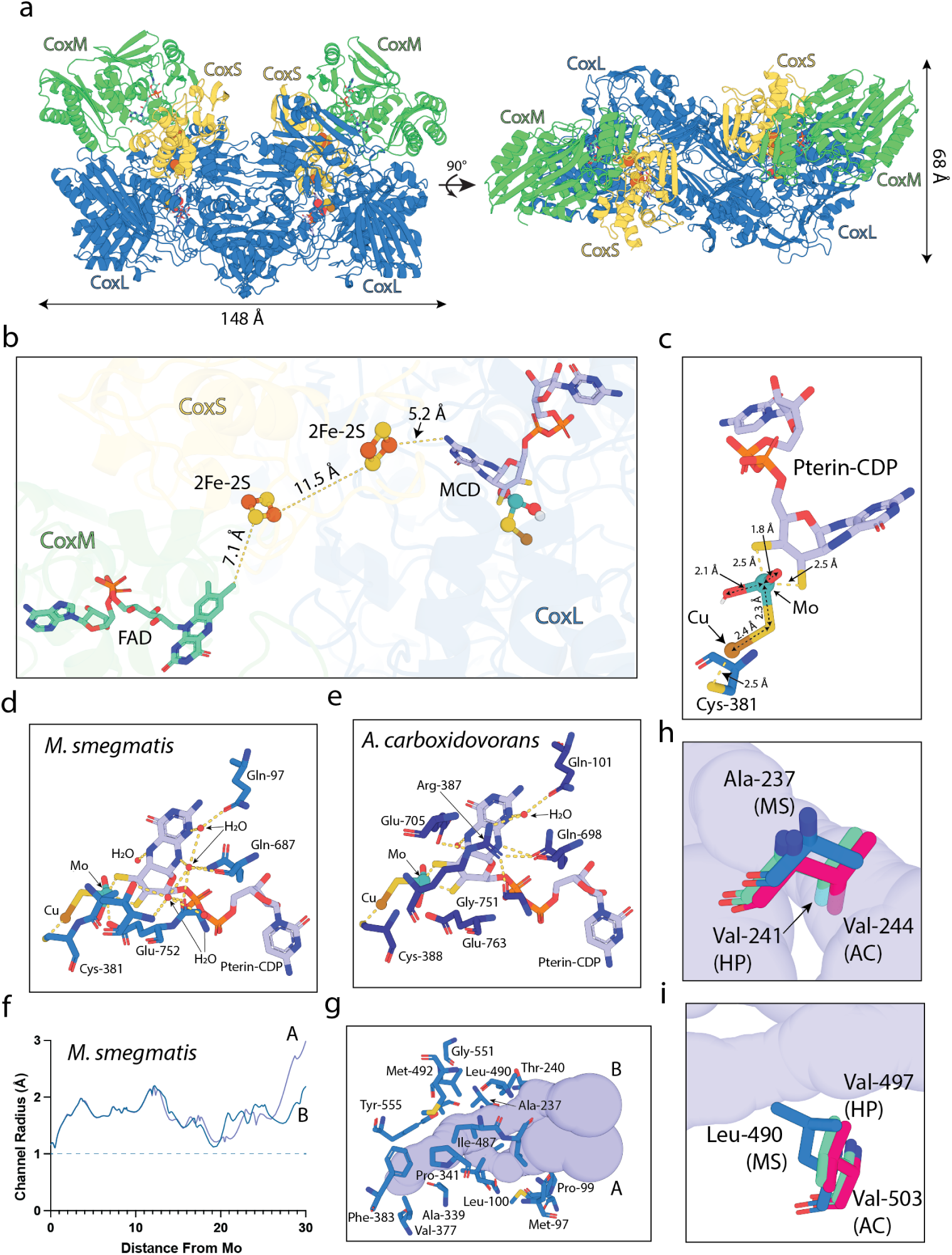
The cryoEM structure of Mo-CODH_Ms_ shows basis of atmospheric CO oxidation. **(a)** A cartoon representation of the cryoEM structure of Mo-CODH_Ms_ showing the CoxS, CoxM and CoxL units and catalytic and electron transfer cofactors. **(b)** A zoomed view of the catalytic and electron transfer cofactors of one subunit of Mo-CODH_Ms_ showing cofactors are positioned within efficient electron-transfer distance. **(c)** The catalytic cluster of Mo-CODH_Ms_, showing bond distance, which indicates the Mo ion is in a reduced (IV) state. **(d)** The coordination environment of the catalytic CuMCD complex in Mo-CODH_Ms_. **(e)** The coordination environment of the catalytic CuMCD complex in Mo-CODH_Ac_. **(f)** A plot showing the radius of the active site gas channel from Mo-CODH_Ms_ relative to the distance from the catalytic Mo ion. **(g)** A zoomed view of the active site gas channel from Mo-CODH_Ms_. **(h)** A zoomed view of Ala-237 from Mo-CODH_Ms_ and the corresponding valine residues from Mo-CODH_Ac_ and CODH_Hp,_ alanine at this position opens the second gas channel in Mo-CODH_Ms._ **(i)** A zoomed view of Leu-490 from Mo-CODH_Ms_ and the corresponding valine residues from Mo-CODH_Ac_ and Mo-CODH_Hp,_ leucine at this position narrows the second gas channel in Mo-CODH_Ms._

The coordination environment of the CuMCD cofactor in Mo-CODH_Ms_ differs markedly from previously structurally characterised Mo-CODHs, including the well-studied Mo-CODH_Ac_ ^16^. In Mo-CODH_Ms_, a conserved arginine (Arg-387 in Mo-CODH_Ac_) is substituted for alanine (Ala-380 in Mo-CODH_Ms_). This leads to significant remodelling of the environment around the CuMCD cofactor in Mo-CODH_Ms_ with three additional highly-coordinated water molecules forming an interaction network that substitutes for the loss of the arginine sidechain (Figure 2d,e; Figure S4b; Movie S3). In the other Mo-CODH to be structurally characterised from *Hydrogenophaga pseudoflava* (Mo-CODH_Hp_) this arginine is post-translationally modified with a hydroxyl group at the C^γ^, suggesting that the specific chemical environment at this position plays a role in tuning the catalytic cofactor ^22^. Phylogenetic analysis of CoxL subunits indicates that arginine is conserved at this position in all Mo-CODH groups, except for the actinobacterial clade that contains Mo-CODH_Ms_, where alanine is universally present (Figure S4c; Supplemental data 1). This substitution may help tune actinobacterial Mo-CODHs for the high-affinity CO oxidation observed in Mo-CODH_Ms_, to support persistence during nutrient deprivation.

The active site of Mo-CODH is in the interior of the CoxL subunit and is accessed by CO via hydrophobic gas channels. These channels have not been previously analysed in Mo-CODH, so we utilised MOLEonline to map them in Mo-CODH_Ms,_ Mo-CODH_Hp,_ and Mo-CODH_Ac_ (Supplemental data 2) ^38^. Interestingly, while both Mo-CODH_Hp_ and Mo-CODH_Ac_ have a single gas channel providing access to the active site, Mo-CODH_Ms_ has two gas channels (Figure 2f,g; Figure S5a-e). Both the gas channels of Mo-CODH_Ms_ are significantly narrower than the gas channel of Mo-CODH_Hp_ and Mo-CODH_Ac_, with bottleneck radii of 1.1 and 1.2 Å, vs 1.4 Å for CODH_Hp_ and 1.6 Å for Mo-CODH_Ac_ (Figure 2f; Figure S5b,d). Substitution of alanine for valine at position 237 of CoxL of Mo-CODH_Ms_ opens the second gas channel, while a number of other mutations constrict the Mo-CODH_Ms_ channels (Figure 2h,g,i; Figure S5f,g). While the more constricted gas channels of Mo-CODH_Ms_ do not appear to drastically restrict the rate of turn over for the enzyme (*k*_cat_ 54.2 s^-1^ vs 93.3 s^-1^ for Mo-CODH_Ac_), they may play a role in modulating the affinity of the Mo-CODH_Ms_ by more effectively capturing substrate CO molecules.

### CoxG has a hydrophobic cavity that binds menaquinone

We next sought to test the hypothesis that Mo-CODH_Ms_ transfers electrons to the aerobic respiratory chain by associating with the membrane-bound protein CoxG. This is consistent with the observations that while isolated Mo-CODH_Ms_ is a soluble enzyme, it does associate with the cytoplasmic membrane in cells, and like Mo-CODH_Ac_, it directly reduces quinone analogues and thus can likely reduce respiratory quinones (Figure 1a) ^4,29^. CoxG is predicted to be comprised of two domains, a C-terminal membrane anchoring helix and an N-terminal soluble cytosolic domain (CoxG_NT_), which are connected by a flexible linker (Figure 3a; Figure S6) ^18,26^.

**Figure 3.**
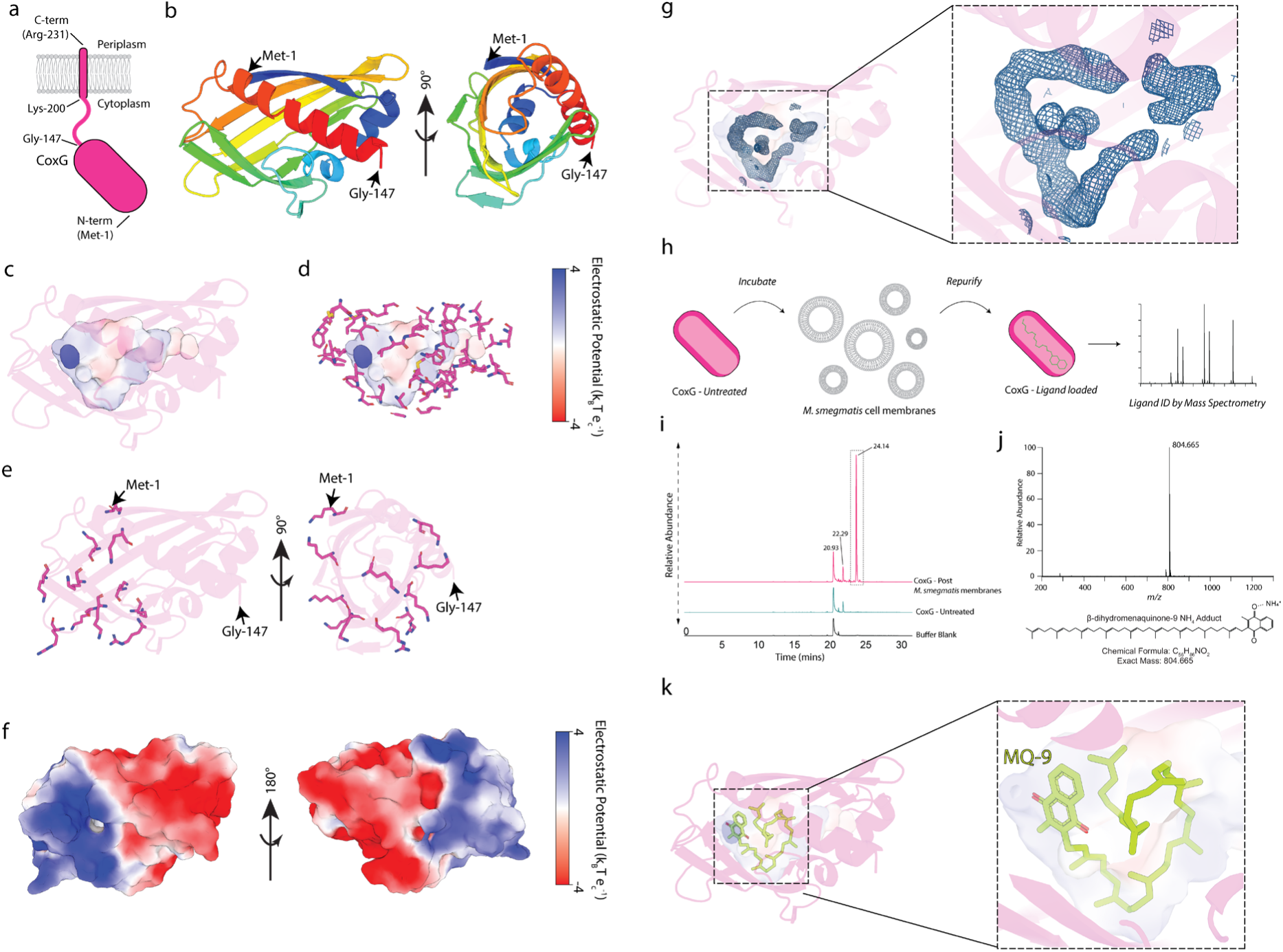
CoxG carries menaquinone-9 within its hydrophobic cavity. **(a)** Schematic of CoxG showing its membrane-anchored C-terminus (CoxG_CT_), cytosolic linker and cytosolic N-terminus (CoxG_NT_). **(b)** A cartoon representation of the X-ray crystal structure of CoxG_NT_ showing the β-sheet and two α-helices that form the SRPBCC fold, with rainbow colouring from N-terminus (blue) to C-terminus (red) **(c)** A surface rendering of the inner hydrophobic cavity of CoxG_NT_. The electrostatic potential of the cavity is shown by blue, white and red colouring. **(d)** A stick representation of the hydrophobic residues that line the inner cavity of CoxG_NT_. **(e)** A stick representation of key positively charged residues present on the N-terminal side of CoxG_NT_. **(f)** An electrostatic surface potential of CoxG_NT_ mapped onto the surface rendering of the structure. Positively charged regions are blue and negatively charged regions are red, with neutral charge in white. **(g)** A composite omit map of electron density corresponding to a lipid-like molecule within the hydrophobic cavity of the crystal structure of CoxG_NT_ contoured at 1 σ. **(h)** A schematic of the experiment performed to test CoxG_NT_ mediated extraction of menaquinone-9 from *M. smegmatis* membranes. **(i)** Base peak chromatograms (m/z = 750 – 2000) for the high-performance liquid chromatography-mass spectrometry (HPLC-MS) analysis of Folch extracts from CoxG_NT_. A substantial peak at 24.14 min is uniquely observed in the *M. smegmatis* membrane-incubated sample and the corresponding positive mode mass spectrum is shown in **(j).** This shows a single dominant ion at m/z = 804.665, consistent with the ammonium adduct of β-dihydromenaquinone-9, m/z = 804.665. **(k)** Molecular docking of MQ9 within the CoxG_NT_ internal cavity, showing the highest-ranked solution.

To gain insight into the functional role of CoxG_NT_, we recombinantly expressed and purified the domain in *E. coli*, and then determined its structure using X-ray crystallography (Table S2). Analysis of the structure revealed that CoxG_NT_ belongs to the SRPBCC superfamily, which is characterised by a seven-stranded antiparallel β-sheet and two α-helices that enclose a large hydrophobic cavity (Figure 3b,c,d; Movie S4). This internal cavity defines the function of the SRPBCC family, which is to bind and traffic diverse hydrophobic molecules through hydrophilic environments ^39,40^. This suggests that, in addition to facilitating the membrane association of Mo-CODH, CoxG may extract a hydrophobic ligand from the membrane. The N-terminal face of CoxG_NT_, which contains an opening to the internal hydrophobic cavity, has a number of lysine residues that create a positively charged region, whereas the C-terminal face that connects to the membrane linker is negatively charged (Figures 3e,f; Movie S2). This positively charged region may interact with the negatively charged head groups of membrane phospholipids, facilitating ligand extraction. In agreement with this, electron density for a lipid-like molecule is present in the internal cavity of the CoxG_NT_ crystal structure (Figure 3g). The identity of this molecule could not be unambiguously determined from the electron density, so we analysed the small molecules associated with purified CoxG_NT_ using mass spectrometry. This analysis detected ubiquinone-8 (UQ8), the major *E. coli* respiratory quinone under aerobic conditions, associated with CoxG_NT_ among a number of other unidentified molecules (Figure S7a,b) ^41,42^. Given the highly hydrophobic character of UQ8, it is likely that it is bound within the CoxG_NT_ cavity, rather than being present in the bulk solvent of the sample. Consistent with this, the electron density present in the CoxG_NT_ cavity is compatible with modelling UQ8, although not at full occupancy (Figure S7c). This observation led us to hypothesize that CoxG shuttles quinone between the membrane and Mo-CODH_Ms_, facilitating its reduction with electrons from atmospheric CO oxidation. The low occupancy of UQ8 in CoxG_NT_ producedin *E. coli* suggests that it is not its preferred ligand, which is consistent with *M. smegmatis* utilising dihydromenaquinone-9 (DH-MQ9) as its major respiratory quinone ^43,44^.

To test the ability of CoxG_NT_ to bind DH-MQ9, we attempted to express it in *M. smegmatis*, where it has the opportunity to extract the ligand from cell membranes. However, it was highly toxic when overexpressed in its native host, possibly due to CoxG_NT_ sequestering cellular DH-MQ9, and purified protein could not be obtained. As an alternative, we incubated CoxG_NT_ produced in *E. coli* with total membranes from *M. smegmatis,* before repurifying it and identifying extracted ligands by mass spectrometry (Figure 3h). Strikingly, the only additional molecule detected in CoxG_NT_ post-incubation was DH-MQ9, which was present at approximately 34% occupancy (Figure 3i,j). Consistent with this, mass spectrometry demonstrated that CoxG_NT_ also bound purified MQ9, despite its sparing solubility in aqueous solution (Figure S7d,e). This indicates that CoxG from *M. smegmatis* selectively binds both DH-MQ9 and MQ9. To model the CoxG-MQ9 complex, we performed docking simulations, which consistently placed MQ9 within the internal cavity of CoxG_NT_, with the highest-ranked solution placing the redox active head group in proximity to the opening of the internal cavity (Figure 3k, Movie S5, Movie S6, Supplemental Data 3). Taken together, both the structure of CoxG_NT_ and its ability to specifically extract MQ9 from membranes support a role in extracting and trafficking respiratory quinone to Mo-CODH for reduction.

### CoxG is essential for CO oxidation in *M. smegmatis* cells and interacts with Mo-CODH

Considering our hypothesis that CoxG transports MQ9 to Mo-CODH_Ms_ for reduction, we hypothesised that deletion of CoxG would decouple it from the respiratory chain and prevent oxidation of CO. Consistent with this, *M. smegmatis ΔcoxG* did not oxidise CO, while as previously reported, the *M. smegmatis* mc^2^155 wildtype strain consumed CO to below atmospheric levels (Figure 4a) ^4^. Complementation of the *ΔcoxG* strain with a plasmid containing *coxG* restored CO consumption to near WT levels (Figure 4a). Although CoxG appears to be essential for CO oxidation in cells, CO oxidation by Mo-CODH_Ms_ was still detectable in cell lysates from the *ΔcoxG* strain, when using the electron acceptor NBT to stain for CO activity (Figure 4b). Mo-CODH_Ms_ activity was lower than wildtype in both the *ΔcoxG* strain and complemented strain (Figure 4b). This is consistent with previous reports in *A. carboxidovorans* where the loss of CoxG led to a 50% reduction in Mo-CODH_Ac_ activity, and could be due to several factors including altered gene expression patterns resulting in a reduction in Mo-CODH expression or loss of activity due to reductive inactivation in the absence of electron transfer via CoxG ^29^.

**Figure 4.**
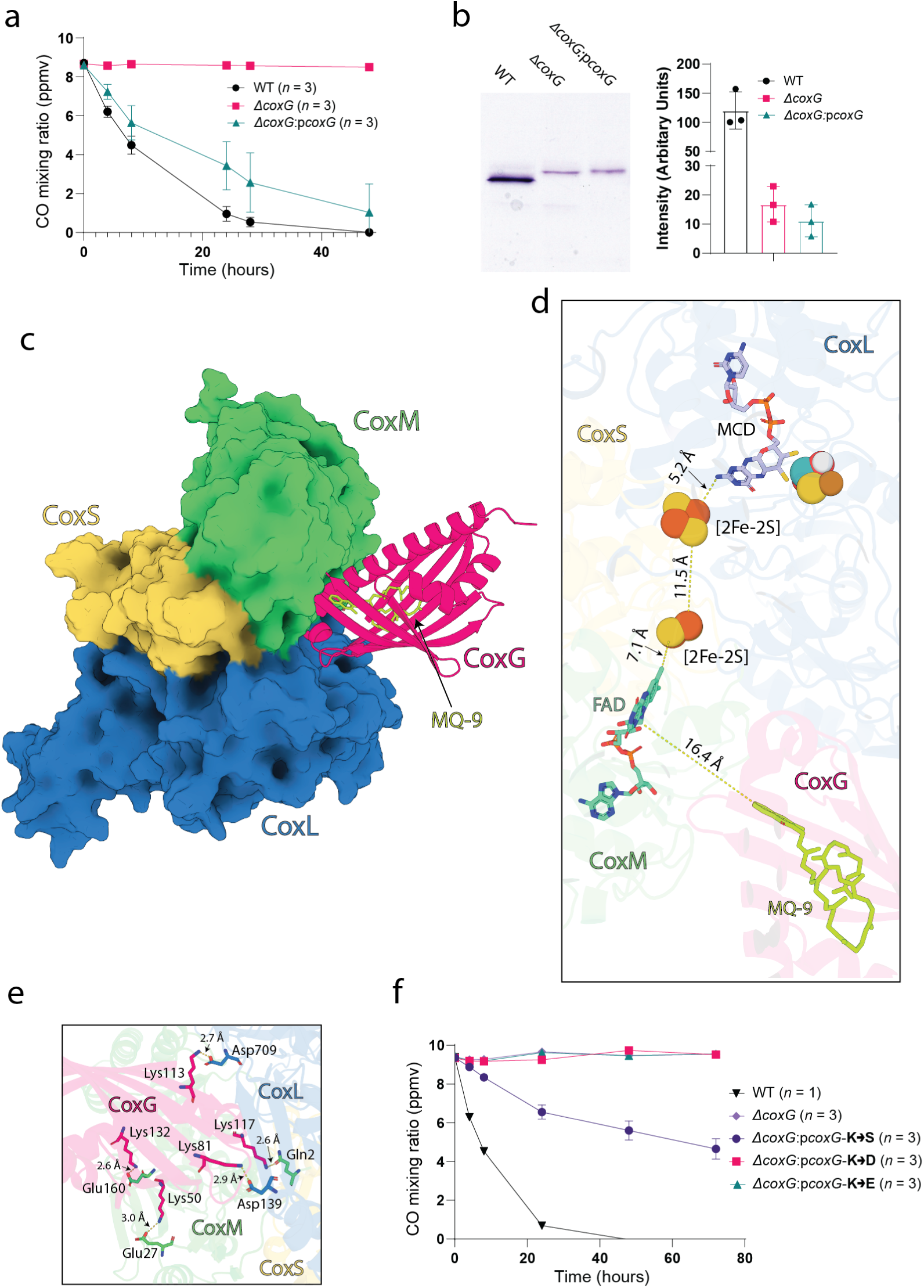
CoxG is predicted to interact with Mo-CODH_Ms_ and facilitate CO oxidation in *M. smegmatis*. **(a)** Gas chromatography analysis of CO in headspace of sealed vials containing the *M. smegmatis* strains WT, *ΔcoxG* or *ΔcoxG:pcoxG*. WT *M. smegmatis* can oxidise CO to below atmospheric levels, however, the *ΔcoxG* strain cannot oxidise CO. Complementation of CoxG on a plasmid restores CO oxidation to near WT levels. **(b)** An NBT-stained Native-PAGE of Mo-CODH_Ms_ from cell lysates of the WT, *ΔcoxG* or *ΔcoxG:pcoxG M. smegmatis* strains. Densitometric analysis normalised to WT staining intensity shows the *ΔcoxG* and *ΔcoxG:pCoxG* strains produce active Mo-CODH_Ms_. **(c)** AlphaFold2 multimer model of the Mo-CODH_Ms_ and CoxG_NT_ (pink) showing a predicted interaction at the interface between CoxM (green) and CoxL (blue). The redox-active head group of MQ9 docked in CoxG_NT_ is positioned towards the Mo-CODH_Ms_ interface. **(d)** The positioning of the CuMCD, [2Fe-2S] and FAD group within Mo-CODH_Ms,_ with distances the electrons transit between each group listed. The predicted interaction between CoxG_NT_ and Mo-CODH_Ms_ positions MQ9 ∼16 Å from the FAD group of Mo-CODH_Ms._ **(e)** CoxG lysine residues that mediate interactions with corresponding negatively charged and polar residues at the CoxG-Mo-CODH_Ms_ interface. **(f)** Gas chromatography analysis of CO headspace of sealed vials containing the *M. smegmatis* strains WT, *ΔcoxG* or *ΔcoxG:pcoxG-***K**→**S***, ΔcoxG:pcoxG-***K**→**D***, ΔcoxG:pcoxG-***K**→**E**. The complementation strains have mutations to the CoxG interface, with the five critical lysines (from (d)) mutated to either serine (K→S), aspartate (K→D) or glutamate (K→E). The WT *M. smegmatis* strain oxidises CO to below atmospheric levels and the *ΔcoxG* strain cannot oxidise CO. The K→D and K→E mutants cannot oxidise CO and the K→S retains some CO oxidation activity, though not to WT levels. In **a**, **b,** and **f** data are mean ± s.d.

To provide DH-MQ9 to Mo-CODH_Ms_ for reduction, CoxG must interact with the enzyme so that electron transfer can occur from the FAD group to bound DH-MQ9. Due to the small quantities of Mo-CODH_Ms_ we can purify, traditional methods for characterising protein-protein interactions were not feasible. As such, we utilised AlphaFold2 multimer to assess complex formation between a single CoxSML subunit of Mo-CODH_Ms_ and CoxG ^45,46^. AlphaFold2 consistently predicts a complex between CoxG and primarily the CoxL and CoxM subunits of CoxLMS (Figure 4c, Supplemental Data 4, Table S3). In this complex, the N-terminal end of CoxG, which contains the opening to the hydrophobic cavity, interacts with Mo-CODH_Ms_ in proximity to the electron-donating FAD group of CoxM. When docked MQ9 is included in this model its redox-active head group is 16 Å away from FAD (Figure 4d). In this position, conformational changes and structural dynamics could allow DH-MQ9 to migrate into the Mo-CODH_Ms_ cavity, which is also hydrophobic, and directly accept electrons from FAD (Figure S8a). The interaction interface is highly charged, containing five lysine residues from CoxG interacting with two glutamate, two aspartate and one asparagine residues from Mo-CODH_Ms_ (Figure 4e, Movie S7). The predominance of lysine residues on the CoxSML interacting interface of CoxG likely stems from its need to interact with the phospholipid headgroups of the cell membrane, in order to extract DH-MQ9 (Figure 3h,i,j). Considering the membrane interacting interface of CoxG must also interact with CoxSML to deliver quinone for reduction by FAD, these positively charged residues must be neutralised by the negatively charged residues observed on the Mo-CODH_Ms_ side of the interface (Table S4). This provides evidence that the surface chemistry of CoxG satisfies interactions with both the cell membrane and its CoxSML binding partner.

To validate the functional importance of the five lysines present at the interaction interface of CoxG in *M. smegmatis*, we generated three variants in which all lysines were mutated to serine (CoxG-K→S), aspartate (CoxG-K→D) or glutamate (CoxG-K→E). Genes encoding these CoxG variants were used to complement the *M. smegmatis ΔcoxG* strain, and consumption of CO was measured by gas chromatography. While the strains expressing CoxG-K→D and CoxG-K→E variants were unable to consume CO, the strain expressing CoxG-K→S regained the ability to oxidise CO, at a significantly reduced rate (Figure 4f). This result is consistent with the CoxG-K→D and CoxG-K→E variants being unable to interact with the lipid membrane and/or CoxSML due to charge repulsion, while the more minor change to polar serine residues impairs but does not totally abolish CoxG function.

### CoxG interactions with Mo-CODH are conserved throughout the enzyme family

To further validate the AlphaFold2 predicted complex between CoxG and Mo-CODH_Ms_, through comparative genomics, we assessed how widespread the presence of *coxG* genes are within the Mo-CODH gene clusters of 30 diverse microbes (Table S5). *coxG* is present in the Mo-CODH gene clusters of 24 of these bacteria and archaea, spanning all five phylogenetic groups of the enzyme. Moreover, in four of the cases where *coxG* was not identified, a homologue is present elsewhere in the genome, which may facilitate quinone transport to Mo-CODH. The only two cases where a *coxG* homologue could not be identified in the Mo-CODH-containing genome were *Sulfolobus islandicus* and *Ca.* Acidianus copahuensis, two archaeal extremophiles, which may have evolved an alternative means of transferring electrons from Mo-CODH ^47,48^. The synteny of *coxG* relative to the *coxSML* structural subunits varies between homologues, and in *Bradyrhizobium japonicum* and *Alkalilimnicola ehrlichii* CoxG is present as a fusion protein with CoxL, further strengthening the functional link between these proteins (Figure 5a). CoxG homologues are also associated with genes encoding non-Mo-CODH members of the xanthine oxidase family (XOF), including an uncharacterised homologue present in *Mycobacterium smegmatis*, where it is positioned between genes encoding the CoxL and CoxM subunits (Figure 5a).

**Figure 5.**
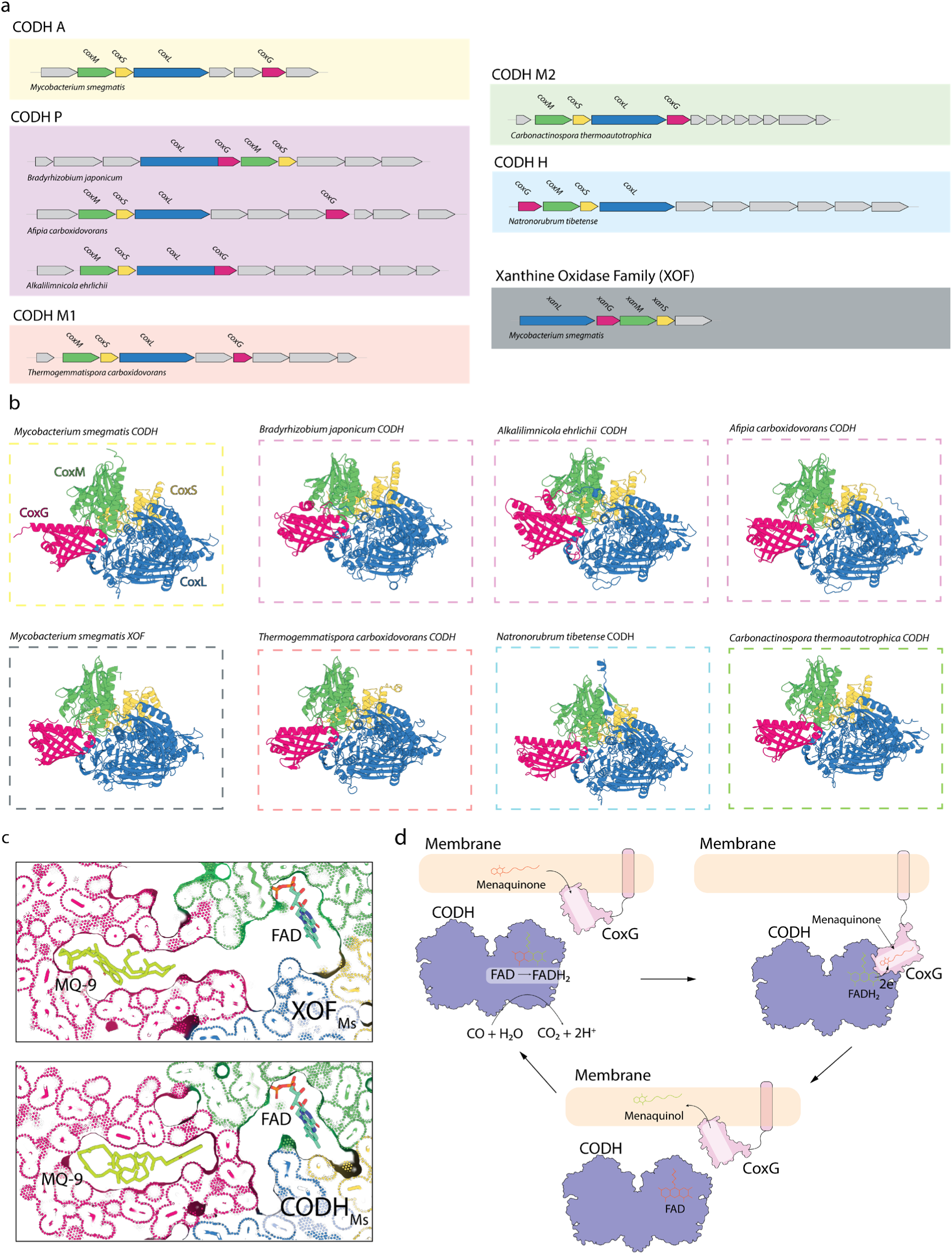
Analysis of the Mo-CODH and CoxG relationship across bacterial species. **(a)** The organisation of Mo-CODH/XOF operons from *M. smegmatis* and six representative microbes. The *coxG* gene (pink) is consistently present in the same gene cluster as the enzymatic subunits of Mo-CODH (*coxM* (green), *coxL* (blue) and *coxS* (yellow)), although with varying synteny and is sometimes fused with *coxL*. **(b)** AlphaFold2 multimer models showing the predicted CoxG and Mo-CODH/XOF complex from the 8 representative gene clusters. All the models predict an analogous complex between CoxG and Mo-CODH/XOF. **(c)** A cut-away dot and surface view of the AlphaFold2 models of Mo-CODH_Ms_ and XOF_Ms_ with the FAD group placed into the CoxM subunit and CoxG docked with MQ9. MQ9 and FAD are contained within a single enclosed cavity in the XOF model (top), whereas in the Mo-CODH_Ms_-CoxG model (bottom), they are enclosed in separate cavities. **(d)** A proposed model for CO oxidation in *M. smegmatis*. Mo-CODH_Ms_ oxidises CO to CO_2_ and the membrane-tethered CoxG extracts MQ9 from the cell membrane and delivers it to Mo-CODH_Ms._ MQ9 is reduced to menaquinol by the Mo-CODH_Ms_ FADH_2_ group and delivered back to the membrane by CoxG, to provide electrons to the terminal oxidases.

To assess whether CoxG also forms a complex with these Mo-CODH homologues, we performed AlphaFold2 multimer modelling with the CoxSML and CoxG subunits of eight representative microbes. In each case, CoxG is predicted to form a complex with the CoxSML subunits analogous to that observed for Mo-CODH_Ms_ (Figure 5b, Figure S8b, Supplemental Data 4). AlphaFold2 predicted the formation of this complex in each Mo-CODH homologue and the *M. smegmatis* XOF, despite an average amino acid sequence identity of 26.4% for the CoxG homologues and 42.6% and 54.6% for the CoxM and CoxL subunits that form the interaction interface (Table S6). The consistency of the AlphaFold2 prediction of this complex across homologues that share limited sequence identity provides strong evidence of its authenticity. When the model of the Mo-CODH_Ms_-CoxG is appended with the CoxM FAD cofactor from the cryoEM structure and CoxG MQ9 from our docking analysis, these cofactors are confined to isolated chambers (Figure 5c), indicating that conformational changes must occur upon CoxG binding to facilitate migration of menaquinone to FAD for reduction. Interestingly, in the *M. smegmatis* XOF model, the interaction between CoxG and CoxSML forms a single enclosed cavity that would allow MQ9 to access the FAD cofactor, suggesting that this model represents a later stage of the substrate transfer process (Figure 5c).

In all Mo-CODH-CoxG complexes, the interactions between CoxG and CoxSML are highly charged, involving multiple salt bridges. Consistent with the Mo-CODH_Ms_-CoxG interface, positively charged sidechains are overwhelmingly present in the CoxG subunit (30 of 33 salt bridges; Table S6). In combination with our other observations, these data support the proposed model for CoxG-mediated quinone transport and its extendibility to most aerobic CO-oxidising microorganisms (Figure 5d).

## Conclusions

In this work, we demonstrate for the first time that an isolated enzyme is capable of oxidising carbon monoxide to sub-atmospheric levels. By determining the high-resolution structure of Mo-CODH_Ms_ we identify key structural differences between this enzyme and low affinity variants that likely contribute to differences in affinity. The coupling of Mo-CODH to the respiratory chain suggested that electrons produced from the oxidation of CO must enter the quinone pool ^4^. However, how Mo-CODH as a soluble enzyme can reduce respiratory quinone was unknown. Based on this work, we propose a model that is conserved across Mo-CODHs from diverse bacteria and archaea, where the lipid binding protein CoxG extracts menaquinone from the membrane before delivering it to the electron acceptor site of soluble Mo-CODH. Menaquinone is then reduced to menaquinol using electrons from atmospheric CO before CoxG returns it to the membrane (Figure 5d). Menaquinol is then oxidised by the terminal-oxidases to generate proton motive force for ATP synthesis and other cellular processes ^49,50^. To our knowledge, this mechanism of electron transfer to menaquinone is unprecedented and mechanistically unique. Moreover, the presence of CoxG in association with diverse Mo-CODHs and other members of the xanthine oxidase family indicates this is a widespread mechanism for coupling soluble enzymes to the respiratory chain. While mechanistically distinct, there are parallels between this system and quinone reduction by Complex I and the recently described [NiFe]-hydrogenase Huc ^43,51^. In both these cases quinone is extracted from the membrane and delivered to its site of reduction via a hydrophobic tunnel ^43,51^. Conversely, our observation that CoxG acts as a quinone shuttle provides a new mechanism of quinone reduction outside the membrane, which evolved convergently to that of Huc and Complex I, and expands our understanding of how electrons enter the respiratory chain.

## Supplemental Figures

**Figure S1.**
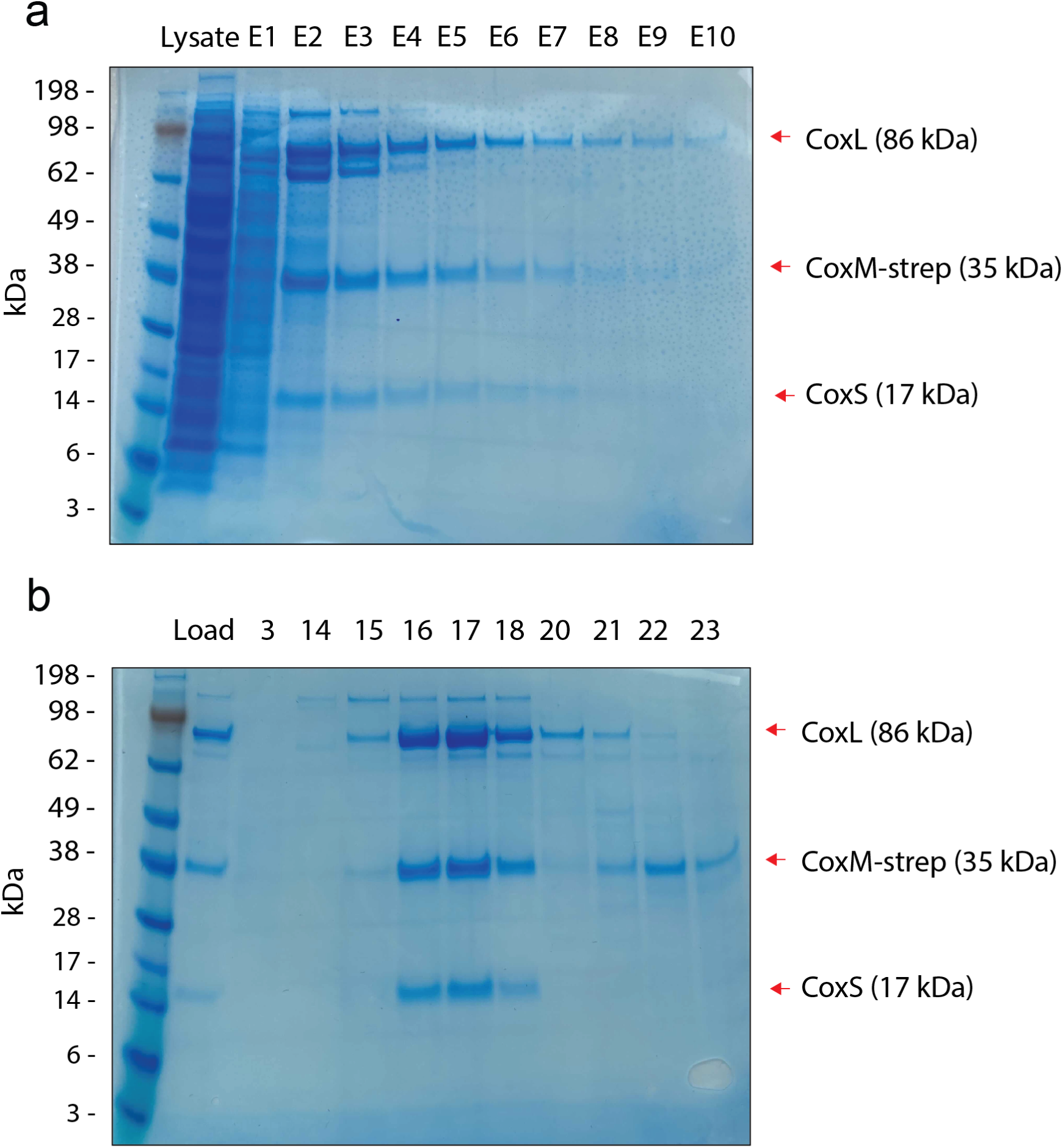
Native purification of Mo-CODH_Ms_. **(a)** SDS-PAGE of Mo-CODH_Ms_ after strep-trap isolation. **(b)** SDS-PAGE of Mo-CODH_Ms_ after SEC, demonstrating the final purity of the sample.

**Figure S2.**
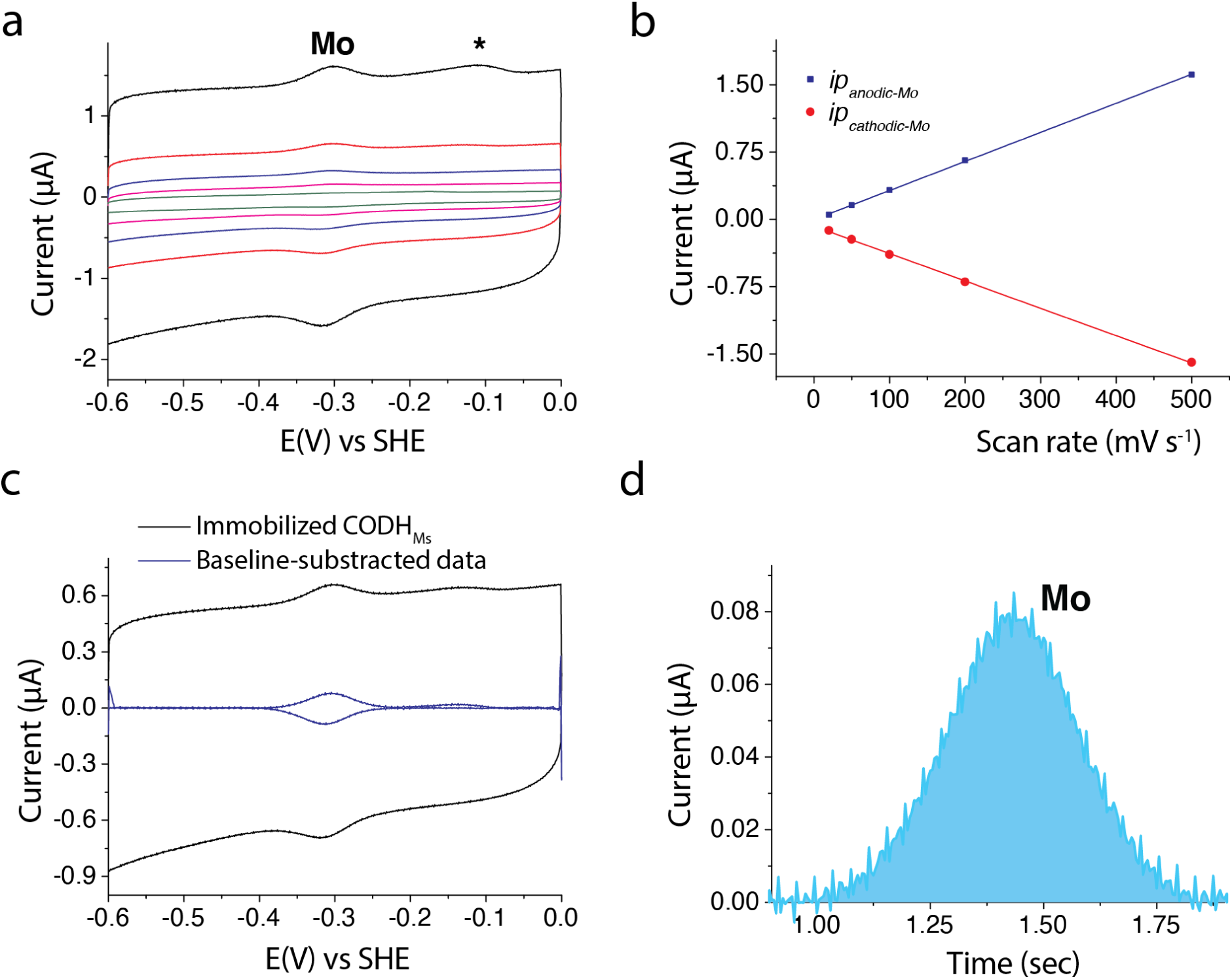
Estimation of loading of Mo-CODH_Ms_ in PFE experiments. **(a)** Scan rate dependent cyclic voltammogram of the surface confined CODH enzyme on a PIGE electrode (black: 500 mV s^−1^, red: 200 mV s^−1^, blue: 100 mV s^−1^, pink: 50 mV s^−1^; green: 20 mV s^−1^)*, T* = 21 °C, pH 8.0. Scan rate dependent anodic and cathodic peak currents (*ip_anodic_*, *ip_cathodic_*) for the Mo processes extracted from panel a. **(c)** DC voltammogram of the surface confined Mo-CODH_Ms_ (black trace), along with the baseline-subtracted data (blue trace) (scan rate = 200 mV s^-1^). **(d)** The background-subtracted oxidation peak associated with the Mo^VI/IV^ process used to obtain the loading of Mo-CODH_Ms_. The potential axis was converted to time using the known scan rate.

**Figure S3.**
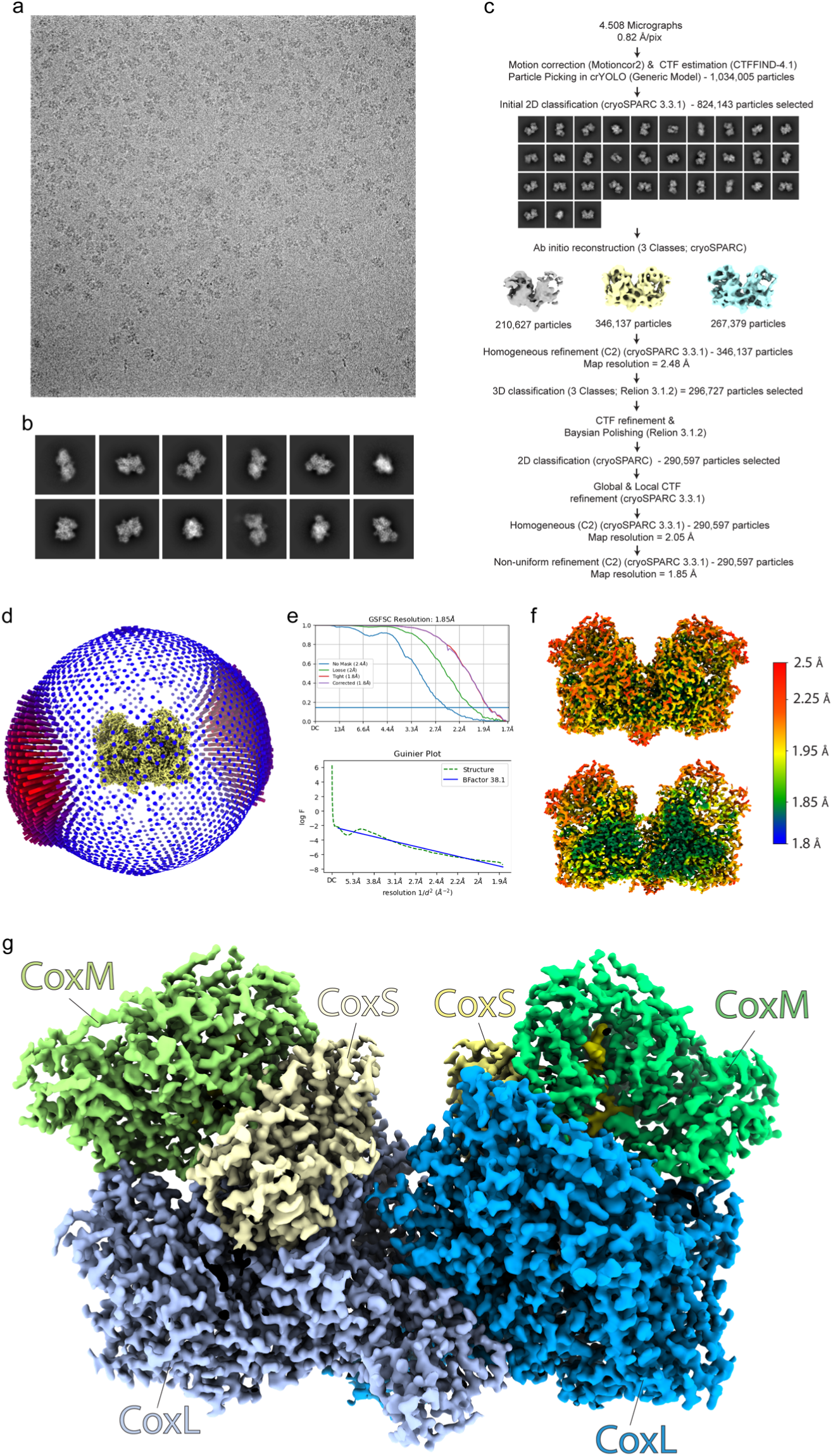
CryoEM data collection, processing, and validation. **(a)** A representative motion corrected cryoEM image of Mo-CODH_Ms_. **(b)** Representative 2D-class averages of Mo-CODH_Ms_ The data processing scheme applied to achieve the 1.85 Å Mo-CODH_Ms_ reconstruction. **(d)** A spherical representation of the relative distribution of particles in the Mo-CODH_Ms_ reconstruction. **(e)** Gold-standard Fourier shell correlation (FSC) curves calculated from two independently refined half-maps indicate an overall resolution of 1.85 Å at FSC = 0.143, and Guinier plot indicates a sharpening B-factor of 38.1. **(f)** A local resolution map of the Mo-CODH_Ms_ oligomer, showing a resolution range of ∼1.8 to 2.5 Å, indicative of limited interparticle motion. **(g)** The final cryoEM density map of Mo-CODH_Ms_ colour coded to show the position of each subunit.

**Figure S4.**
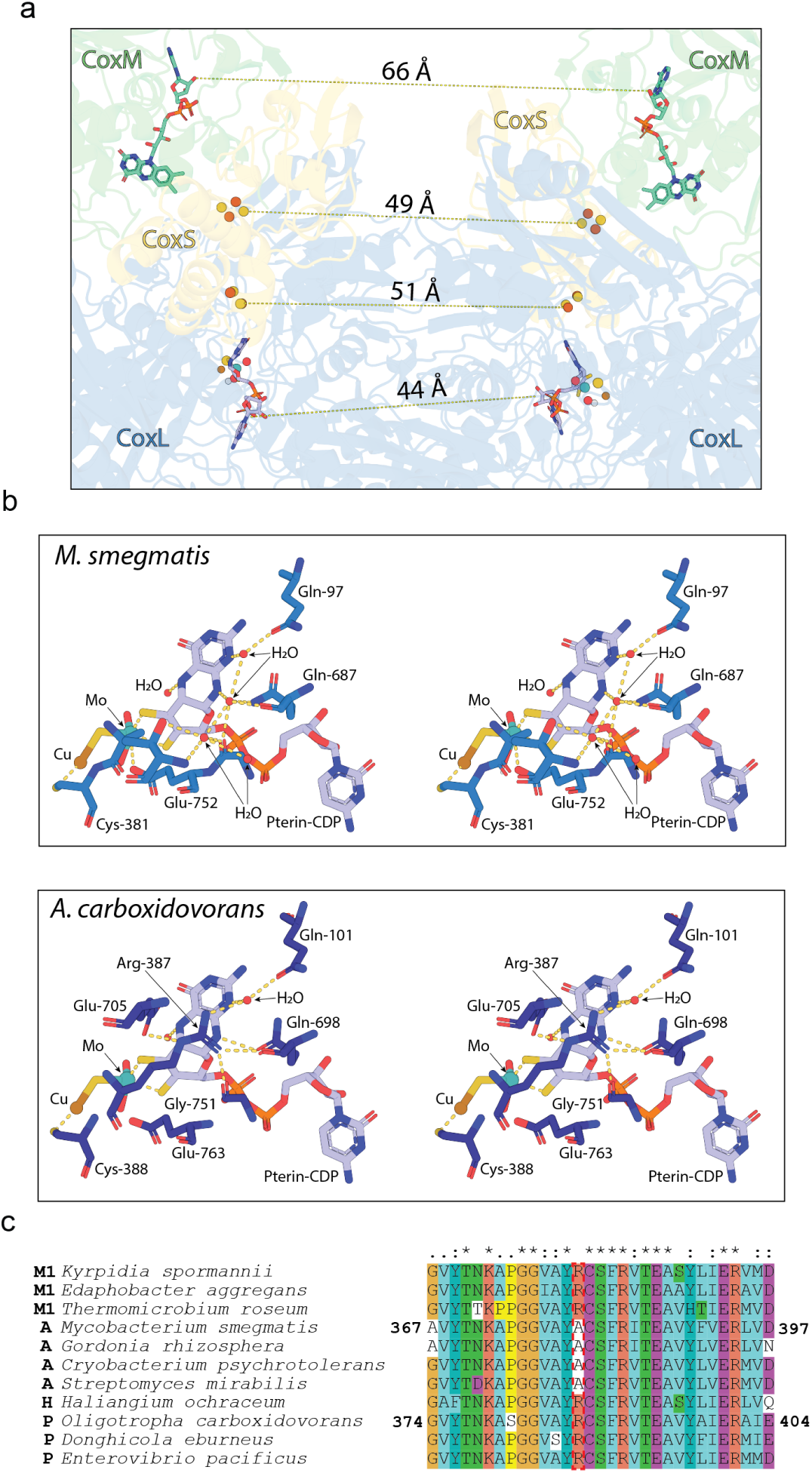
Mo-CODH_Ms_ structural, catalytic and sequence analysis. **(a)** A zoomed view of the structure of Mo-CODH_Ms_ showing the electron transferring cofactors are outside of efficient electron transfer distance. **(b)** A cross-eyed stereo view of the coordination environment of the catalytic CuMCD complex in Mo-CODH_Ms_ (top), the catalytic CuMCD complex in Mo-CODH_Ac_ (bottom). **(c)** A sequence alignment of the catalytic cluster coordinating region of CoxL from representative Mo-CODHs showing the substitution of arginine (R) for alanine (A) in actinobacterial group enzymes. M1 = Mixed group 1, A = Actinobacterial group, H = Haloarchaeal group, P = Proteobacterial group.

**Figure S5.**
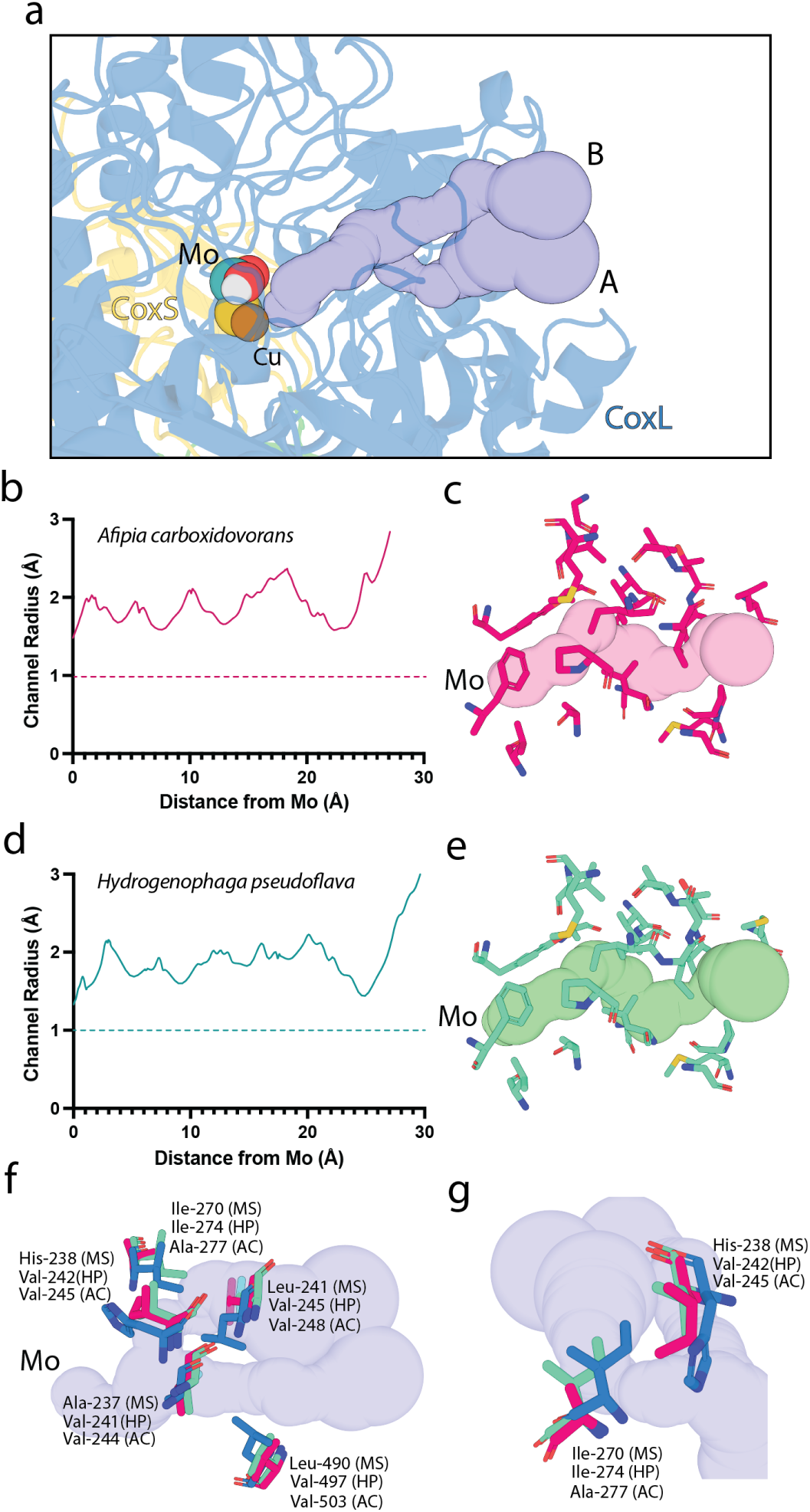
Comparison of substrate transporting gas channels from Mo-CODHs. **(a)** A zoomed cartoon view of Mo-CODH_MS_ showing the two gas channels that mediate gas exchange with the active site. **(b)** A plot showing the radius of the active site gas channel from Mo-CODH_Ac_ relative to the distance from the catalytic Mo ion. **(c)** A zoomed view of the active site gas channel from Mo-CODH_Ac_. **(d)** A plot showing the radius of the active site gas channel from Mo-CODH_Hp_ relative to the distance from the catalytic Mo ion. **(e)** A zoomed view of the active site gas channel from Mo-CODH_Hp_. **(f), (g)** A comparison of divergent residues that line the active site gas channels of Mo-CODH_Ms_ (HP), Mo-CODH_Ac_ (AC), and Mo-CODH_Hp_ (HP), which reshape the gas channels of Mo-CODH_Ms._

**Figure S6.**
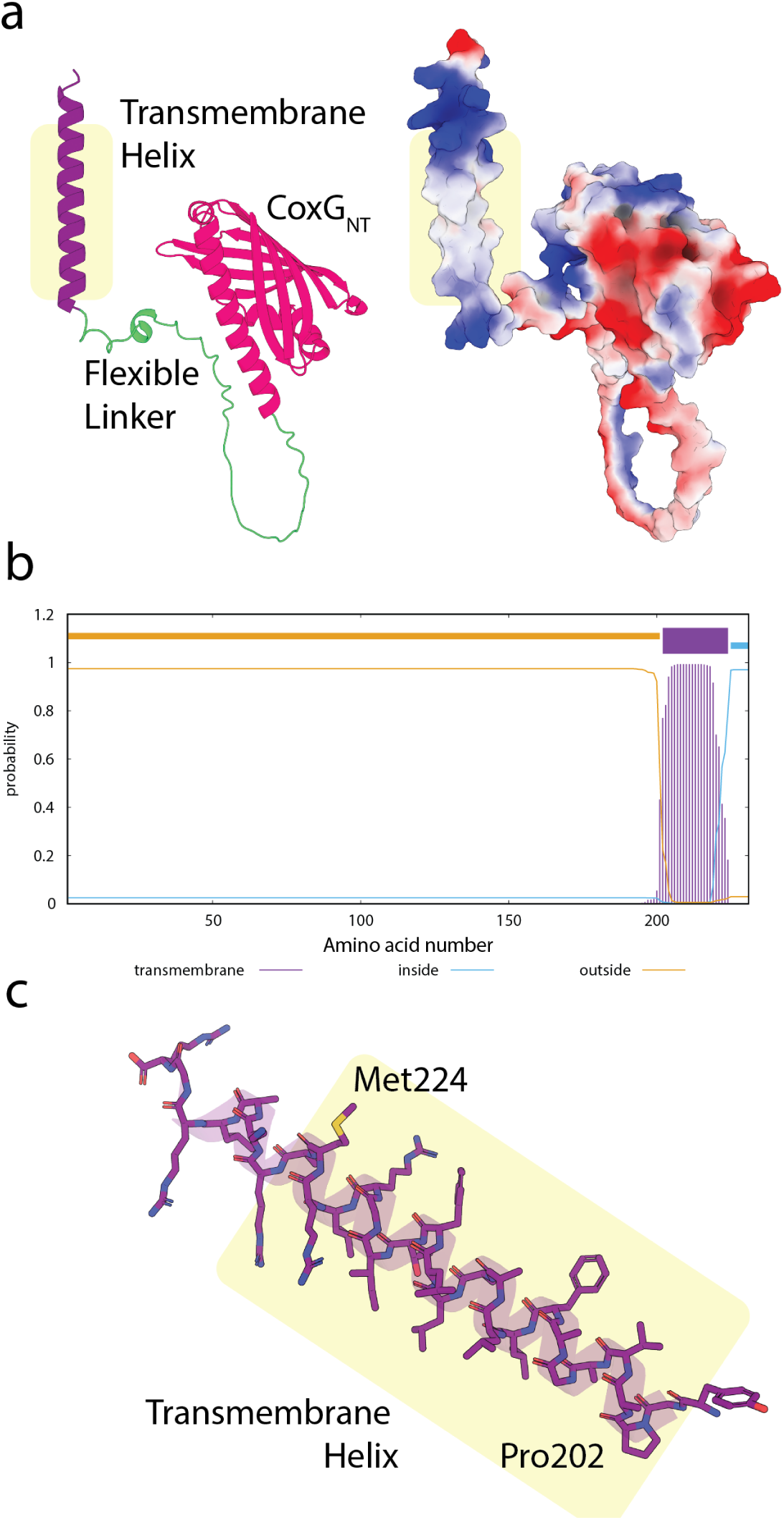
CoxG contains a C-terminal transmembrane helix. **(a)** An AlphaFold2 model of full length CoxG from *M. smegmatis*, showing the three functional domains as a cartoon view (left) and the electrostatic surface (right). The transmembrane region predicted by TMHMM - 2.0 is highlighted by a yellow box. **(b)** A plot showing the prediction of membrane spanning helices in CoxG by TMHMM - 2.0 ^52^. **(c)** A zoomed view of the transmembrane helix of CoxG from *M. smegmatis*.

**Figure S7.**
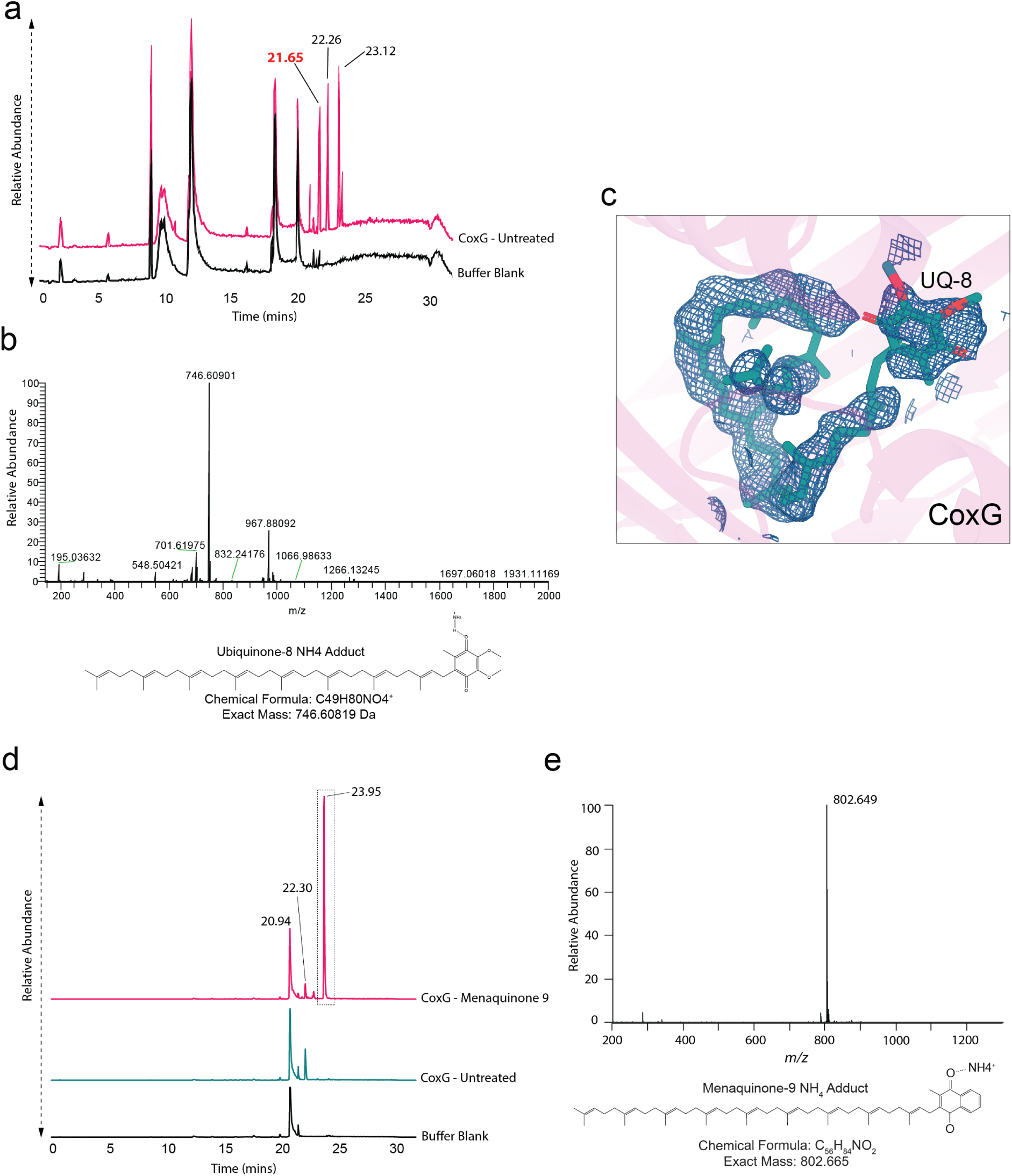
LC-MS identification of CoxG_NT_ associated molecules. **(a)** High performance liquid chromatography–mass spectrometry (LC–MS) analysis of the Folch extract from purified CoxG_NT_. The base peak chromatogram (left) shows a peak at 21.65 min, corresponding to an ion at m/z = 746.609 **(b)**, which is consistent with the ammonium adduct of ubiquinone-8, m/z =746.608. **(c)** The composite omit map of electron density within the hydrophobic cavity of the crystal structure of CoxG_NT_ contoured at 1 σ, fitted with ubiquinone-8. **(d)** High-performance liquid chromatography-mass spectrometry (LC-MS) analysis of Folch extracts from CoxG_NT_, with a substantial peak at 23.95 min in the menaquinone-9 incubated sample. **(e)** The mass spectrum of the species eluting at 23.95 min, which has a m/z = 802.649, consistent with the ammonium adduct of menaquinone-9, m/z = 802.665.

**Figure S8.**
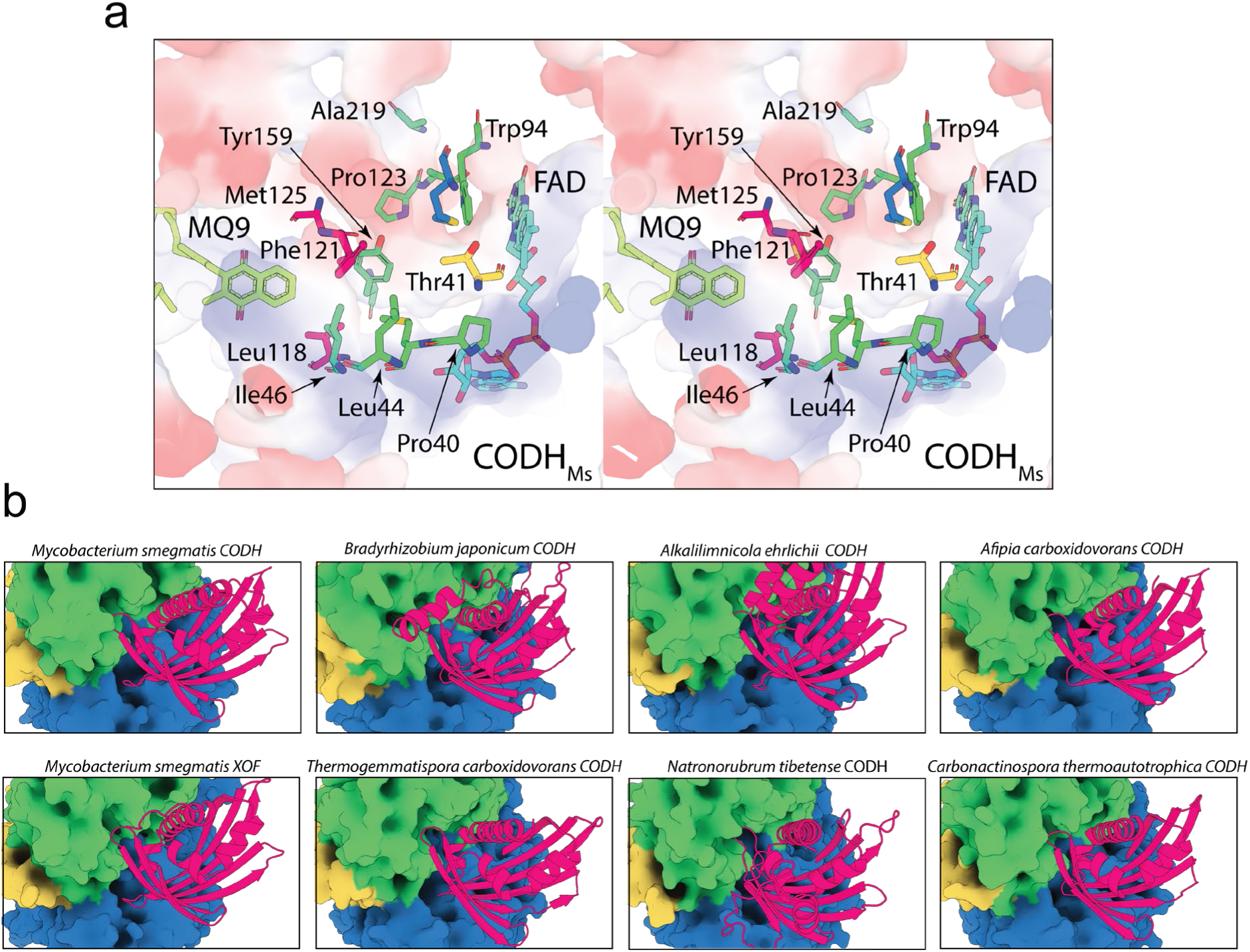
Analysis of the AlphaFold2 modelled CoxG-CoxSML interface. **(a)** A stereo view of the CoxG-Mo-CODH_MS_ complex, showing a composite view of the external electrostatic surface of the complex and a stick view of hydrophobic residues that form this surface. The position of MQ9 and FAD are also shown as sticks. **(b)** A zoomed in view of the CoxG interacting surface of Mo-CODH in the AlphaFold2 models from various species.

## Supplemental Notes

**Supplemental Note 1. Protein film electrochemistry of Mo-CODH_Ms_**

Cyclic voltammograms were recorded on a Mo-CODH_Ms_ film in electrolyte solutions at pH 8.0 under a N_2_, CO or CO_2_ atmosphere (Figure 1d, e). Under the N_2_ atmosphere, a well-defined chemically reversible process was observed with a mid-point potential of −0.305 V (Figure 1d). In addition, two oxidation peaks were found at the potential of −0.120 and +0.185 V (Figure 1d). Under CO atmosphere, a sigmoidal-shaped catalytic oxidation process emerged at the potential region where the chemically reversible process took place (Figure 1e, red trace). These results suggest the process at −0.305 V is associated with the redox transformation of the molybdenum active center, which involves two electrons (Mo^VI/IV^)^34^. The reversible potential associated with the Mo^VI/IV^ process is < 0.3 V more positive than the equilibrium potential of the CO_2_/CO reaction (−0.58 V vs SHE at pH 8.0^31^), demonstrating the effectiveness of Mo-CODH_MS_ in catalysing this process. Our electrochemical data are consistent with the previous report by Bernhardt et al. on the *A. carboxidovorans* Mo-CODH enzyme who found the Mo^V/IV^ and Mo^VI/V^ processes with reversible potentials of −0.357 V and −0.165 V respectively, in an electrolyte solution containing 50 mM HEPES (pH = 7.2) based on EPR measurements undertaken at liquid N_2_ temperatures.^53^ In that study, the authors also found processes at more positive potential region which were assigned to the FeS^I^, FeS^II^ clusters, and FAD cofactors, which likely correspond to the oxidation peaks at −0.120 V and +0.185 V observed in our data. Two closely spaced Mo^VI/V^ and Mo^V/IV^ processes have often been found for the same type of enzymes ^54,55^.

The sigmoidal shape of the voltammogram obtained under a CO atmosphere suggests that the voltammetric process is kinetically controlled. Therefore, a turnover frequency (*k_cat_*) of 54.2 s^-1^ can be obtained from the catalytic current plateau (*i*_cat_) based on the following equation using an enzyme loading (*N)* value of 1.7×10^-13^ moles (see below for calculation) estimated from the voltammogram.

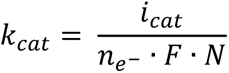

Where *n_e_^−^* and *F* refer to the number of electrons involved in the reaction (two in this case) and Faraday’s constant, respectively. This *k_cat_* value is close to that of 93.3 s^-1^ associated with the Mo-CODH from *A. carboxidovorans* (pH 7.2, 25 °C)^32^. No catalytic reduction of CO_2_ was observed under CO_2_-saturated conditions (see Figure 1d; blue line). This is expected since Mo-CODH are limited to CO oxidation.^56,57^ The redox potential of the enzyme is much more positive than that of CO_2_/CO, making the CO_2_ reduction reaction thermodynamically unfavorable.

Enzyme loading was determined from DC voltammograms of immobilized Mo-CODH_Ms_ at pH 8.0 (Figure S2a). The linearity of the peak current versus scan rate indicates that the enzyme is successfully confined on the electrode surface (Figure 2b). The amount of charges (*Q*) associated with the oxidation Mo-CODH was estimated from the peak area of the background subtracted voltammogram (Figure S2c, d). Then, a *N* value of 1.7×10^-13^ moles was calculated using Faraday’s law (*N* = *Q*/(*n_e_-F*)), which corresponded to a surface coverage of 1.9×10^−1^^2^ mol/cm^2^, typical for enzyme monolayers on electrodes^58^.

## Materials and methods

### Bacterial culturing

*M. smegmatis* MC^2^155 WT, CoxM-strep, *ΔCoxG, ΔcoxG:pcoxG* and *ΔcoxG:pcoxG* mutant strains were grown from frozen glycerol stocks on lysogeny broth agar plates supplemented with 0.05% (w/v) Tween80 (LBT) for 3-4 days at 37°C (Table S7). Starter cultures were inoculated with colonies from LBT plates and were grown in LBT media at 37°C and shaking at 200 rpm overnight. Broth cultures were inoculated from turbid starter cultures to OD_600_ = 0.01 and grown in Hartman’s de Bont (HdB) minimal media supplemented with 0.05 % (v/v) glycerol and 0.05% (w/v) tyloxapol and incubated at 37 °C with shaking at 150 rpm. Cells were harvested at 3 days post-OD_max_ (peak Mo-CODH_MS_ expression).

### Mutant construction

*M. smegmatis* MC^2^155 chromosomal mutants were generated using allelic exchange mutagenesis as previously described ^43,59^. A twin-Strep II tag was inserted at the C-terminus of the *coxM* gene (MSMEG_0744) of one strain, and a knock-out of the *coxG* gene was performed in another. 500 bp upstream and downstream of the *coxM* C-terminus were fused to a twin-Strep II tag (5’GGCGGTTCGGGCTGGTCCCACCCCCAGTTCGAAAAGGGTGGGGGCTCCGGTGGCGGGTCGGGTGGGTCCGCCTGGTCGCACCCGCAGTTCGAGAAG 3’) in a 1111 bp fragment synthesised by Genscript. Two ∼500bp fragments upstream and downstream of the *coxG* gene were fused to create a deletion construct of 1011 bp and synthesised by Genscript. The fragments were cloned into the SpeI site of the mycobacterial shuttle plasmid pX33 and transformed into *M. smegmatis* mc^2^155 via electroporation ^59^. To allow permissive temperature–sensitive vector replication, transformants were incubated on LBT gentamycin plates at 28°C until colonies were visible (7 days). The resultant colonies were subcultured in LBT gentamycin broth at 40°C for 3-5 days and then diluted from broth onto fresh LBT gentamycin plates and incubated at 37°C for 3–5 days to facilitate integration of the recombinant plasmid, via flanking regions, into the chromosome. The second recombination event was facilitated by subculturing gentamycin-resistant colonies in LBT supplemented with 10% sucrose (w/v) for 3–5 days, followed by dilution onto LBT agar plates supplemented with 10% sucrose (w/v) and incubation at 37°C for 3–5 days. Gentamycin-sensitive colonies were subsequently screened by PCR to distinguish WT revertants from mutants. The primers used for screening are listed in Table S7.

The CoxG WT complementation plasmid was constructed by PCR of *coxG* from *M. smegmatis* mc^2^155 gDNA followed by ligation into the pMV261 complementation vector using the BamHI and EcoRI restriction sites (primer sequences listed in Table S7). CoxG complementation mutants were synthesised as gene fragments by Twist Bioscience and sub-cloned into pMV261 using the restriction sites BamHI and EcoRI.

### Purification of Mo-CODH_Ms_

*M. smegmatis* mc^2^155 CoxM-strep was cultured in 8 L of HdB until 3 days post max OD_600_. Pellets were harvested via centrifugation (30 min at 5,000 × *g*) and stored at −20 °C. Pellets from 16-24 L batches of culture were thawed, pooled and resuspended in ice-cold lysis buffer (150 mM NaCl, 50 mM Tris pH 8.0, 1 × cOmplete EDTA-free protease inhibitor (Roche), 1 mg/mL DNase and lysozyme) using a Dounce glass homogenizer. Cells were lysed via two passages through a cell disrupter (Emulsiflex C-5) and cell debris was removed via centrifugation at 30,000 × *g* for 30 minutes. The clarified cell lysate was loaded onto a 1 mL StrepTrap XT column (Cytiva) and washed with ∼100 mL chromatography buffer (150 mM NaCl, 50 mM Tris pH 8.0). Protein was eluted with elution buffer (150 mM NaCl, 50 mM Tris pH 8.0, 50 mM biotin) in 10 × 1 mL fractions. Biotin was dissolved in the buffer by adjusting the pH back to pH 8.0 with 5 M NaOH after mixing. Elution fractions were analysed by SDS-PAGE and fractions containing Mo-CODH_Ms_ were pooled and concentrated to a volume of ∼500 µL using a 4 mL 30K MWCO centrifugal concentrator (Amicon). Concentrated fractions were loaded onto a Superose 6 Increase 10/300 GL column (Cytiva) and eluted in 0.5 mL fractions using chromatography buffer and analysed via SDS-PAGE. Fractions containing Mo-CODH_Ms_ were pooled and concentrated to ∼25 µL at 2-4 mg/mL using a 0.5 mL 30K MWCO centrifugal concentrator (Amicon) and stored at −80 °C.

### Expression and purification of CoxG

For recombinant expression, p29b-CoxG (pCoxGtrunc) was transformed *E. coli* (DE3) C41 cells were cultured in terrific broth (per litre: 12 g tryptone, 24 g yeast extract, 61.3 g K_2_HPO_4_, 11.55 g KH_2_PO_4_, 10 g glycerol, with 100 µg/ml ampicillin for selection) as previously described^61^ (Table S7). Cells were grown at 37°C until they reached an OD_600_ of 1.0 and were then induced using 0.3 mM IPTG (isopropyl-β-d-thiogalactopyranoside), followed by further growth for 14 h at 25°C. Cells were harvested by centrifugation, lysed using a cell disruptor (Emulsiflex) in Ni-binding buffer (50 mM Tris, 500 mM NaCl, 20 mM imidazole [pH 7.9]) plus 0.1 mg/ml lysozyme, 0.05 mg/ml DNase I, and cOmplete protease cocktail inhibitor tablets (Roche). The resulting lysate was clarified by centrifugation and applied to Ni-agarose resin, followed by washing with 10× column volumes of Ni-binding buffer, and elution of bound proteins with a step gradient of Ni-gradient buffer (50 mM Tris, 500 mM NaCl, 500 mM imidazole [pH 7.9]) of 5, 10, 25, and 50%. Eluted fractions containing recombinant protein were pooled and applied to a 26/600 Superdex S200 SEC column equilibrated in SEC buffer (50 mM Tris, 200 mM NaCl [pH 7.9]). The recombinant protein was then pooled, concentrated to 60 mg/ml, snap-frozen, and stored at −80°C.

### PAGE analysis, activity staining and western blotting

For SDS- and native-PAGE, samples were run on Bolt 4–12% SDS-polyacrylamide and native PAGE 4-16% gels (Invitrogen) respectively, according to the manufacturer’s instructions. Gels were visualized by AcquaStain Protein Gel Stain (Bulldog), or nitrotetrazolium blue (NBT) activity staining. For NBT activity staining, gels were incubated in 50 mM Tris, 150 mM NaCl pH 8.0 supplemented with 500 μM NBT in an anaerobic jar amended with 100% CO for 24 hours at RT. Bands exhibiting Mo-CODH_Ms_ activity were identified based on the purple colour of reduced NBT.

### Gas chromatography

For culture-based CO consumption assays, *M. smegmatis* wild-type (WT), *coxG* KO and *ΔcoxG:pCoxG* cultures were grown in 30 mL Hartmans de Bont (HdB) medium amended with 0.05% (v/v) glycerol and 0.05% (w/v) tyloxapol in 120 mL glass serum vials, with incubation at 37 °C and shaking at 150 rpm. When cultures reached stationary phase (3 days post-ODmax) the vials were sealed with butyl rubber stoppers and amended with 10 ppm CO gas. Consumption was monitored over 48 hours at regular time intervals by injecting 2 mL of headspace into a pulsed discharge helium ionisation detector (model TGA-6791-W-4U-2, Valco Instruments Company Inc.) as previously described ^12^.

For enzymatic CO consumption assays, 5 mL reaction buffer (50 mM Tris, 150 mM NaCl, pH 8.0, 0.3 mg/mL BSA, 50 µM methylene blue or 200 µM menadione) was purged with N_2_ (5 min, intermittent shaking) in 120 mL glass serum vials sealed with a butyl rubber stopper and magnetic stirrer bar. Purged vials were amended with purified Mo-CODH_Ms_ to 8 nM or 5 nM, or vehicle control (reaction buffer without Mo-CODH_Ms_), using a 250 µL gas-tight syringe, and then immediately amended with 1 ppm or 10 ppm CO (10 ppm cal gas, BOC, CCS405162D2) and incubated at room temperate with gentle stirring. Reactions for the condition were prepared in triplicate. Seven time points were taken over ∼70 hours, with t0 taken 30 s after starting incubation. Headspace CO concentration was measured by injecting 2 mL of headspace into a pulsed discharge helium ionisation detector (model TGA-6791-W-4U-2, Valco Instruments Company Inc.) as previously described ^12^.

For Mo-CODH_Ms_ kinetics, 5 mL of reaction buffer (50 mM Tris, 150 mM NaCl, pH 8.0, 0.3 mg/mL BSA, 50 µM methylene blue) was added to a 120 mL glass serum vials sealed with a butyl rubber stopper and purged with N_2_. CO was added to the vial headspace to a final concentration of 1, 5, 50, 200, 500 or 1000 ppm (1, 3.5, 40, 250, 500, 850 nM dissolved CO in the buffer). Either 1 nM, 2 nM or 5 nM of Mo-CODH_Ms_ was added to the reaction (lower CO concentrations required more Mo-CODH_Ms_ for consumption) for it to begin. All reaction vessels were in triplicate. CO consumption was measured as described, using a pulsed discharge helium ionisation detector, by injecting 2 mL of headspace at 8 time points over 8 hours. The rate of CO oxidation was calculated for each CO concentration as nmole.hr^-1^ CO consumed by 1 nM of Mo-CODH_Ms_. This was plotted as dissolved CO (nM) vs. rate (nmole.hr^-1^) and using the Michaelis-Menten non-linear fit function on GraphPad Prism, *V*_max_ and *K*_m_ were calculated.

### Protein film electrochemistry of Mo-CODH_Ms_

A CH Instruments 760E potentiostat was used to conduct the direct current (DC) voltammetric experiments. A standard three-electrode system in a single compartment cell was used. The set-up consisted of a Ag/AgCl (sat’d KCl) reference electrode, a platinum wire as the counter electrode, and a bare custom-built paraffin-impregnated graphite rod (PIGE)^58^ (electroactive surface area, *A* = 0.09 cm^2^) or a Mo-CODH_MS_ enzyme modified PIGE as the working electrode. To fabricate the Mo-CODH_MS_ enzyme modified PIGE, the Mo-CODH_Ms_ enzyme was immobilized on the electrode by placing the electrode vertically and carefully pipetting 2 µL of a 0.32 mg/mL stock solution of the purified enzyme (stored in 150 mM NaCl, 50 mM Tris buffer aqueous solution at pH 8.0). The electrode was placed under a gentle flow of N_2_ for 5 minutes (until dry), and the surface was cleaned with a small amount of distilled water followed by drying with N_2_. All voltammetric experiments were conducted in an electrolyte solution containing 150 mM NaCl and 50 mM Tris buffer at pH 8.0. Prior to the voltammetric experiments, the electrolyte solution was degassed with N_2_, CO or CO_2_ for at least 10 minutes. During the voltammetric experiments, a positive gas pressure was applied to ensure that oxygen was not re-dissolved in the solution. All voltammograms were recorded at 21±1 **°**C. The potential scale of all voltammograms reported here was converted into the standard hydrogen electrode (SHE) scale knowing the potential of the Ag/AgCl (sat’d KCl) reference electrode is +0.204 V vs SHE. at 20 **°**C.^60^

### CryoEM imaging of Mo-CODH_Ms_

Samples (3 μl) were applied onto a glow-discharged UltrAuFoil grid (Quantifoil GmbH) and were flash frozen in liquid ethane using the Vitrobot mark IV (Thermo Fisher Scientific) set at 100% humidity and 4 °C for the prep chamber. Data were collected on a G1 Titan Krios microscope (Thermo Fisher Scientific) with S-FEG as electron source operated at an accelerating voltage of 300 kV. A C2 aperture of 50 μm was used and no objective aperture was used. Data was collected at a nominal magnification of 105 K in nanoprobe EFTEM mode. Gatan K3 direct electron detector positioned post a Gatan BioQuantum energy filter was operated in a zero-energy-loss mode using a slit width of 10 eV to acquire dose fractionated images of the Mo-CODH_Ms_ complex. One dataset was collected, composed of 4,508 movies. Movies were recorded in hardware-binned mode yielding a physical pixel size of 0.82 Å pixel^−1^ with a dose rate of 8.5 e− pixel^−1^ s^−1^. An exposure time of 6 s yielded a total dose of 66.0 e− Å^−2^, which was further fractionated into 60 subframes. Automated data collection was performed using EPU (Thermo Fisher Scientific) with periodic centring of zero loss peak. A defocus range was set between −1.5 and −0.5 μm.

### CryoEM data processing and analysis

Micrographs from all datasets were motion-corrected using UCSF Motioncor and dose-weighted averages had their CTF parameters estimated using CTFFIND 4.1.8, implemented using Relion 3.1.2 ^61^. Particle coordinates were determined by crYOLO 1.7.6 using a model trained on manual particles picked from 20 micrographs ^62^. Unbinned particles were extracted from micrographs using Relion 3.1.2, before being imported into cryoSPARC 3.3.1 for initial 2D classification to remove bad particles, followed by ab initio model generation and 3D refinement^63^. Refined particles were reimported into Relion 3.1.2 and CTF refinement was performed, followed by Bayesian Polishing^61^. Particles were reimported into cryoSPARC 3.3.1 for final 2D classification to remove residual bad particles, followed by non-uniform 3D consensus refinement to generate final maps at 1.85 Å (FSC = 0.143, gold standard).

### Mo-CODH_Ms_ cryoEM model building and visualization

An initial model of the CoxMSL subunit trimer was generated using AlphaFold2 and docked into one-half of the high-resolution CoxMSL homodimer maps using ChimeraX^67,72^. The model was refined and rebuilt into map density using Coot^64^. CuSMoO_2_, molybdopterin, [2Fe-2S], and FAD cofactors associated with Mo-CODH_Ms_ were downloaded from the PDB:1N5W and fitted and refined into maps using Coot ^16,64^. The model was then refined using real-space refinement within PHENIX ^65^. Once model building was complete the model was symmetry expanded using the Map symmetry tool and waters were added using DOUSE within the PHENIX package^65^. Model quality was validated using MolProbity^66^. Images were generated in Pymol^67^.

### CoxG_NT_ crystallisation, data collection, and structure solution

Purified CoxG was screened for crystallisation conditions using commercially available screens (∼800 conditions). Diffraction data were collected at 100 K at the Australian Synchrotron and processed using the XDS package ^68^. Phases were obtained using an AlphaFold model of CoxG from *M. smegmatis* ^45^. The initial model was built using Phenix ^65^. The CoxG model was improved manually in Coot and refined using Phenix refine^64,65^. Analysis of the CoxG crystal structure was performed using the Phenix and CCP4 packages ^69^. Figures were made using Pymol and ChimeraX ^67,70^.

### Mo-CODH tunnel analysis using MOLEonline

MOLEonline was used to measure the radii of Mo-CODH substrate channels for Mo-CODH_Ac_ (PDB ID: 1N5W), Mo-CODH_Hp_ (PDB ID : 1FFV) and our cryoEM structure of *M. smegmatis* Mo-CODH ^38^. The setting ‘Ignore HETATMS’ was turned off. To prevent the interpretation of artefacts, the settings ‘Probe Radius’, ‘Interior Threshold’ and ‘Origin Radius’ were explored over multiple submissions for each protein complex and the consistency between predictions was monitored, and the substrate channels were inspected using the embedded viewer and in Pymol after export. Cavity data was exported and visualised in GraphPad Prism and Pymol.

### AlphaFold structural modelling

AlphaFold2 modelling was performed using AlphaFold version 2.1.1 implemented on the MASSIVE M3 computing cluster ^45,46^. For the prediction of the CoxG-CoxMSL complex, the corresponding sequence with the flexible C-terminal extension of CoxG removed was provided and modelling was run in multimer mode, with just one molecule of each subunit requested. The five ranked models produced by AlphaFold were compared for consistency with the top-ranked model utilised for further analysis and figure generation. Inter-subunit interfaces were analysed with PISA^71^. Models are provided in **Supplemental data 4**.

### Molecular docking of MQ9 into CoxG_NT_

To determine the ability of CoxG_NT_ to bind MQ9, we utilised Autodock Vina in the Chimera software package ^72,73^. Coordinates for MQ9 were extracted from PDB entry 1DXR. This structure was optimised and refinement restraints were generated using the eLBOW in the Phenix package ^65^. A search box was set encompassing the entire CoxG_NT_ crystal structure and a total of nine binding modes were sought for each docking run, with search exhaustiveness of between eight and 300 and a maximum energy difference of 3 kcal/mol. An identical procedure was performed on an AlphaFold2 model of the CoxG_NT_ from XOF from *M. smegmatis*.

### Mass spectrometric menaquinone detection

Purified CoxG expressed in *E. coli* was incubated with *M. smegmatis* membranes extracted from 8 L of *M. smegmatis* culture, or pure MQ9 suspended in DMSO. To prepare membranes, *M. smegmatis* mc^2^155 was grown in LBT for 3 days at 37 °C and harvested by high-speed centrifugation. Pellets were resuspended in assay buffer (50 mM Tris, 150 mM NaCl pH 8.0, supplemented with plus 0.1 mg/ml lysozyme, 0.05 mg/ml DNase I, and cOmplete protease cocktail inhibitor tablets (Roche)) and lysed using a cell disrupter (Emulsiflex). Cell debris was pelleted by centrifugation (18K RPM, 30 mins) and the clarified lysate was subjected to ultracentrifugation (100,000 ×g, 1 hour). Membranes were resuspended in assay buffer and incubated with 5 mg of CoxG overnight at 4 °C, slowly rotating. The membranes were pelleted through ultracentrifugation again, and CoxG was repurified using Ni-affinity chromatography with a Ni-agarose resin. Following loading onto the column, the resin was washed with 10× column volumes of Ni-binding buffer, and elution of bound proteins with a step gradient of Ni-gradient buffer (50 mM Tris, 150 mM NaCl, 500 mM imidazole [pH 8.0]) of 5, 10, 25, and 50%. For MQ9 incubation, 5 mg (277 nmol) of CoxG in 50 mM Tris, 150 mM NaCl pH 8.0, was mixed with 1 mg (1,273 nmol) of MQ9 suspended in DMSO, in a final volume of 100 µL. MQ9 appeared to be sparingly soluble under these conditions and formed a waxy coating on the reaction tube. The solution was incubated overnight at 20 °C before CoxG was repurified as described for the membrane incubation. Samples corresponding to 12 µg of purified protein, together with a positive control containing MQ9 equivalent to the quantity of ligand assuming complete occupancy (calculated based on the following: m = 12 µg, MW_CoxG_ = 18 kDa, MW_MQ9(II-H2)_ = 786.63 g/mol, n = 12 x 10^-6^/18 x 10^3^ = 0.667 nmoles. As each protein contains a single ligand: n_ligand_ = 0.667 nmoles; m_MQ9_ = 0.667 x 10^-9^ x 786.63 = **524.42 ng**), were prepared for liquid chromatography-mass spectrometry (LC-MS) analysis using a modified Folch extraction. Briefly, a 50 μL solution of purified protein (∼12 μg) was treated with 1000 μL of 2:1 chloroform:methanol v/v after which the mixture was shaken for 10 mins and allowed to stand for a further 50 mins. 200 μL of water was added and the mixture was shaken for 10 mins after which the sample was allowed to stand until the two phases had completely separated. The lower chloroform-rich phase was then transferred to a 2 mL sample vial and the solvent was removed under a stream of nitrogen gas. The resulting residue was reconstituted in 100 μL of 2:1 chloroform:methanol v/v and transferred to a 200 μL sample insert, the solvent was again removed and the sample reconstituted in a 7:3 mixture of LC solvent A:LC solvent B v/v. Samples were analysed using a Dionex RSLC3000 UHPLC (Thermo) coupled to a Q-Exactive Plus Orbitrap MS (Thermo) using a C18 column (Zorbax Eclipse Plus C18 Rapid Resolution HD 2.1 x 100mm 1.8 micron, Agilent) with a binary solvent system; solvent A = 40% isopropanol and solvent B = 98% isopropanol both containing 2 mM formic acid and 8 mM ammonium formate. Linear gradient time-%B as follows: 0 min-0%, 8 min-35%, 16 min-50%, 19 min-80%, 23 min-100%, 28 min-100%, 30 min-0%, 32 min 0%. The flow rate was 250 μL min^-1^, the column temperature 50°C, and the sample injection volume was 10 μL. The mass spectrometer operated at a resolution of 70,000 in polarity switching mode with the following electrospray ionization source conditions: Spray voltage 3.5kV; capillary temperature 300 °C; sheath gas 34; Aux gas 13; sweep gas 1 and probe temp 120 °C. The percentage occupancy of the ligand was estimated by comparing the peak area from the sample and the MQ9 sample.

### Mass spectrometric identification of CoxSML

1 µL of purified Mo-CODH_Ms_ was sent to the Monash Proteomics and Metabolomics Facility for detection. Proteins were reduced with tris(2-carboxyethyl)phosphine (Pierce), alkylated with iodoacetamide (Sigma), and digested with mass spectrometry grade trypsin (Promega). The extracted peptides were analysed by LC-MS/MS on an Ultimate 3000 RSLCnano System (Dionex) coupled to an Orbitrap Fusion Tribrid (ThermoFisher Scientific) mass spectrometer equipped with a nanospray source. The peptides were first loaded and desalted on an Acclaim PepMap trap column (0.1 mm id × 20 mm, 5 μm) and then separated on an Acclaim PepMap analytical column (75 μm id × 50 cm, 2 μm) over a 30 min linear gradient of 4–36% acetonitrile/0.1% formic acid. The Orbitrap Fusion Tribrid was operated in data-dependent acquisition mode with a fixed cycle time of 2 s. The Orbitrap full ms1 scan was set to survey a mass range of 375– 1800 m/z with a resolution of 120,000 at m/z 400, an AGC target of 1 × 10^6^, and a maximum injection time of 110 ms. Individual precursor ions were selected for HCD fragmentation (collision energy 32%) and subsequent fragment ms2 scan were acquired in the Orbitrap using a resolution of 60,000 at m/z 400, an AGC target of 5 × 10^5^, and a maximum injection time of 118 ms. The dynamic exclusion was set at ±10 ppm for 10 s after one occurrence. Raw data were processed using Byonic (ProteinMetrics) against a protein database covering *Mycobacterium smegmatis* mc^2^155. The precursor mass tolerance was set at 20 ppm, and fragment ions were detected at 0.6 Da. Oxidation (M) was set as dynamic modification, carbamidomethyl-(c) as fixed modification. Only peptides and proteins falling below a false discovery rate of 0.01 were reported.

### Phylogenetic analysis of Mo-CODH gene clusters

For sequence analysis of CoxL subunits, 709 amino acid sequences compiled by Cordero et al.^4^, were aligned using the Clustal algorithm ^74^, and the conservation of key residues was assessed manually. For a detailed analysis of the synteny of Mo-CODH gene clusters a representative subset of Mo-CODH sequences presented in Grinter and Greening 2022 was utilised. Out of the 33 Mo-CODHs analysed in this work, 30 available genome sequences were analysed for the presence of CoxG homologues in the Mo-CODH gene cluster or elsewhere in the genome, using the NCBI genome browser. CoxG, CoxS, CoxM and CoxL sequences from six of these Mo-CODH (representing all Mo-CODH groups) and XOF from *M. smegmatis* were utilised to generate AlphaFold2 models of the CoxG-CoxSML complex, as described above.

## Acknowledgements

We acknowledge the use of instruments and assistance at the Monash Ramaciotti Centre for Cryo-Electron Microscopy, a Node of Microscopy Australia. This work was supported by an Australian Research Council (ARC) DECRA Fellowship (DE170100310) (to C.G.), an ARC Discovery Project Grant (DP200103074) (to R.G. and C.G.), a National Health and Medical Research Council Emerging Leader Grant (NHMRC) (APP1178715) (to C.G.), a NHMRC Emerging Leader Grant (APP1197376) (to R.G.), ARC LIEF grants (LE200100045, LE120100090) for the Titan Krios Gatan K3 Camera and for the Titan Krios. We thank David Steer and the Monash Proteomics and Metabolomics Facility for the mass-spec identification service we used in this study.

